# Fine tuning of hormonal signaling is linked to dormancy status in sweet cherry flower buds

**DOI:** 10.1101/423871

**Authors:** Noémie Vimont, Adrian Schwarzenberg, Mirela Domijan, Armel S. L. Donkpegan, Rémi Beauvieux, Loïck le Dantec, Mustapha Arkoun, Frank Jamois, Jean-Claude Yvin, Philip A. Wigge, Elisabeth Dirlewanger, Sandra Cortijo, Bénédicte Wenden

## Abstract

In temperate trees, optimal timing and quality of flowering directly depend on adequate winter dormancy progression, regulated by a combination of chilling and warm temperatures. Physiological, genetic and functional genomic studies have shown that hormones play a key role in bud dormancy establishment, maintenance and release. We combined physiological, transcriptional analyses, quantification of abscisic acid (ABA) and gibberellins (GAs), and modelling to further investigate how these signaling pathways are associated with dormancy progression in the flower buds of two sweet cherry cultivars.

Our results demonstrated that GA-associated pathways have distinct functions and may be differentially related with dormancy. In addition, ABA levels rise at the onset of dormancy, associated with enhanced expression of ABA biosynthesis *PavNCED* genes, and decreased prior to dormancy release. Following the observations that ABA levels are correlated with dormancy depth, we identified *PavUG71B6*, a sweet cherry *UDP-GLYCOSYLTRANSFERASE* gene that up-regulates active catabolism of ABA to ABA-GE and may be associated with low ABA content in the early cultivar. Subsequently, we modelled ABA content and dormancy behavior in three cultivars based on the expression of a small set of genes regulating ABA levels. These results strongly suggest the central role of ABA pathway in the control of dormancy progression and open up new perspectives for the development of molecular-based phenological modelling.

## INTRODUCTION

Perennial plants have evolved strategies that enhance survival under the various environmental stresses they face during their growth and reproductive cycles. Among them, dormancy is a quiescent phase that protects meristematic and reproductive tissues from freezing damage. In temperate trees, the transition from active growth to dormancy is often triggered by decreasing photoperiod and/or temperatures depending on the species (Heide and Prestrud 2005, Rohde et al. 2011, Petterle et al. 2013, Singh et al. 2016). Subsequently, the bud dormancy process relies on the integration of cold and warm temperatures between endodormancy, when buds are unable to resume growth even under favorable conditions, and ecodormancy, when bud development is inhibited by unfavorable conditions until optimal growth temperatures and photoperiod are met (Lang et al. 1987). In the current context of climate change, temperate trees are affected by contradictory effects during the dormancy period and shifts in phenological phases are observed: longer growing season and insufficient chilling during winter, both effects potentially having dramatic impact on growth and production (Vitasse et al. 2011, Atkinson et al. 2013, Jochner et al. 2013). Dormancy progression and control by temperature and photoperiod in perennial plants have been a focus for decades and physiological, genetic and functional genomic studies have shed some light onto the mechanisms underlying dormancy control in deciduous trees and other perennial plants (Cooke et al. 2012, Ríos et al. 2014, Beauvieux et al. 2018). Bud dormancy is controlled by a complex array of signaling pathways that integrate endogenous and environmental cues towards a rest/growth decision. Effort to synthesize the available knowledge and data into modelling approaches have led to the development of phenological models based on the dormancy regulation by temperature and photoperiod (Chuine et al. 2016, Chuine and Régnière 2017). However, process-based models of bud dormancy have not changed substantially since 1990 (Hänninen 1990) and the current predictive models rely on very little information about involved mechanisms. Conceptual models for dormancy progression have been proposed based on interactions between respiratory stresses, ethylene and abscisic acid (ABA), which in turn activate gibberellins (GA)-mediated growth through up-regulation of *FLOWERING LOCUS T* (*FT*) expression and resumption of intercellular transport (Ophir et al. 2009, Rinne et al. 2011).

The major role of hormones in the regulation of bud growth cessation, dormancy and activity resumption has been extensively discussed (e.g. Cooke et al. 2012, Beauvieux et al. 2018, Liu and Sherif 2019). Seed and bud dormancy show common features in terms of hormonal control (Powell 1987, Leida, Conejero, et al. 2012, Wang et al. 2016) and GA and ABA balance is often involved in the integration of internal and external cues to control plant growth (Rodríguez-Gacio et al. 2009, Finkelstein 2013, Shu et al. 2018): GAs promote growth, whereas ABA promotes dormancy. Multiple physiological and transcriptomic studies have indeed proposed a central role for ABA in the repression of bud activity during dormancy. ABA would function as a signal in response to autumn short days and decreasing temperatures to induce dormancy onset (Rohde et al. 2002, Rohde and Bhalerao 2007, Ruttink et al. 2007, Wang et al. 2016, Tuan et al. 2017, Li et al. 2018, Tylewicz et al. 2018). Strong correlation was further shown between ABA and dormancy depth with high ABA levels detected during endodormancy, followed by a decrease in endogenous ABA content during the transition from endodormancy to ecodormancy (Or et al. 2000, Zheng et al. 2015, Wang et al. 2016, Wen et al. 2016, Chmielewski et al. 2017, Li et al. 2018, Zhang et al. 2018, Yamane et al. 2019). Recently, ABA content has been proposed as a determining factor to assess dormancy status in sweet cherry (Chmielewski et al. 2017) and we have shown in a previous study that genes involved in ABA-related pathways were central in the transcriptomic analysis of flower bud dormancy (Vimont et al. 2019). More precisely, recent transcriptomic analyses of genes involved in the precise balance between biosynthesis and catabolism modulating ABA levels have further defined the involvement of ABA in bud dormancy. Indeed, expression patterns for 9-cis epoxycarotenoid dioxygenases (*NCED*), that catalyze the critical step for ABA biosynthesis, and *CYP707A*, encoding cytochrome P450 monooxygenases that inactivate ABA into 8’hydroxy ABA, as well as ABA signaling genes, support ABA involvement in bud dormancy induction and maintenance (Fig. **1a**; Nambara and Marion-Poll 2005, Bai et al. 2013, Zhong et al. 2013, Zhu et al. 2015, Wang et al. 2016, Khalil-Ur-Rehman et al. 2017, Li et al. 2018, Zhang et al. 2018, Zheng et al. 2018, Yu et al. 2020). Similarly, until recently, most of the knowledge gathered on the behavior of the GA pathway during dormancy had been obtained in seeds but reports published in the last years have shed some light on GA regulation throughout bud dormancy in perennial plants. Studies have suggested a major role for GAs in maintaining growth before the induction of dormancy (Junttila and Jensen 1988, Ruttink et al. 2007, Olsen 2010, Eriksson et al. 2015, Singh et al. 2016) and promoting growth during ecodormancy (Wen et al. 2016, Zhang et al. 2018). Interestingly, GA treatments have a controversial effect on dormancy and bud break as shown in various perennial species since GA application may substitute for chilling (Shafer and Monson 1958, Rinne et al. 2011, Zhuang et al. 2013), or have delaying effects on shoot growth and bud break (Hoad 1983, Zheng et al. 2018), suggesting distinct gibberellin functions during dormancy. Although transcriptomic results for *GA2ox, GA3ox* and *GA20ox* vary between studies and therefore suggest complex and distinct functions, general patterns could be identified: expression for GA biosynthesis genes *GA 20-oxidases* (*GA20ox*) and *GA 3-oxidases* (*GA3ox*) decreases during dormancy induction then increases after dormancy release and during ecodormancy while GA deactivation *GA 2-oxidases* (*GA2ox*) genes are up-regulated during endodormancy and inhibited after endodormancy is released (Fig. **1b**; Yamaguchi 2008, Bai et al. 2013, Zhong et al. 2013, Zhu et al. 2015, Wen et al. 2016, Khalil-Ur-Rehman et al. 2017, Zhang et al. 2018, Zheng et al. 2018).

In this study we explored potential hormonal markers of dormancy using a combination of physiological and transcriptomic analyses and a new modelling approach. We have focused on the involvement of GA and ABA pathways in sweet cherry flower bud dormancy. We examined the effect of exogenous GA and ABA on dormancy status and monitored endogenous contents for GAs and ABA and its metabolites, as well as the expression of genes related to ABA and GA metabolism throughout flower bud dormancy for two cultivars with contrasted dormancy release dates. Following our findings on hormonal control of dormancy, we propose a mathematical model that incorporates the effect of key genes on the dynamics of ABA to estimate dormancy status.

## MATERIALS AND METHODS

### Plant material

As previously described (Vimont et al. 2019), samples were collected from three different sweet cherry cultivars (*Prunus avium* L.) having very early, early and late flowering dates (respectively, ‘Cristobalina’, ‘Garnet’ and ‘Regina’). Trees are grown in an orchard located at the Fruit Experimental Unit of INRA in Bourran, South West of France (44° 19′ 56′′ N, 0° 24′ 47′′ E) under standard agricultural practices. During the sampling season (July 2015 to March 2016), a mix of randomly chosen flower buds (equivalent to a 2 mL volume) were sampled at ten time points spanning the entire period of bud development (Fig. **2a**) for phytohormone quantification (for ‘Cristobalina’ and ‘Regina’) and RNA-seq analysis (for the three cultivars). Flower buds were harvested from branches of three (‘Cristobalina’ and ‘Garnet’) or two different trees (‘Regina’). A total of 29, 31 and 21 samples were analyzed for ‘Cristobalina’, ‘Garnet’ and ‘Regina’ respectively. Details are available in Table **S1** (Supplementary file at *Tree Physiology* online). Upon harvesting, buds were flash frozen in liquid nitrogen and stored at - 80°C prior to performing RNA-seq. Average daily temperatures were recorded by an on-site weather station.

In addition, for the exogenous application of hormones, branches were collected from the late flowering sweet cherry cultivar ‘Fertard‘. Trees were grown in an orchard located at the Fruit Experimental Unit of INRA in Toulenne, South West of France (48° 51′ 46′′ N, 2° 17′ 15′′ E) under standard agricultural practices.

We calculated the chill accumulation from September 1^st^ for each sampling date and the dates of dormancy release, using chilling hours (CH), i.e. the sum of hours when temperatures are above 0°C and below 7.2°C (Weinberger 1950), and chill portions (CP, estimated by the Dynamic model; Fishman et al. 1987a, 1987b).

### Measurements of bud break and estimation of the dormancy release date

Measurements for the dormancy stages were performed on randomly chosen branches cut every two weeks from November 16th 2015 to April 4th 2016 for ‘Cristobalina’, ‘Garnet’, ‘Regina’ and ‘Fertard’, and between November 21st 2017 and April 4th 2018 for ‘Fertard‘. Branches were incubated in water pots placed in a growth chamber (25°C, 16h light/ 8h dark, 60-70% humidity). The water was replaced every 3-4 days. After ten days under these forcing conditions, the percentage of bud break, i.e. flower buds at BBCH stage 53 (Fadón et al. 2015), as illustrated in Fig. **S1a** (Supplementary file at *Tree Physiology* online), was recorded. The date of dormancy release was estimated when at least 50% of the flower buds were at the BBCH stage 53 or higher after ten days under forcing conditions.

### Treatments with exogenous hormones and antagonists

To investigate the effects of GA and ABA on dormancy, five branches per modality were randomly harvested from ten ‘Fertard’ dormant trees on January 19th 2016 and January 29th 2018 (Fig. **S1**). The cherry dormant buds were treated with 5 μM GA_3_ (Sigma-Aldrich, ref. 48870), 5 μM GA_4_ (Sigma-Aldrich, ref. G7276), 400 μM ABA (Sigma-Aldrich, ref. A1049), 300 μM paclobutrazol (Sigma-Aldrich, ref. 46046), an inhibitor of the GA pathway, and 5 μM fluridone (Sigma-Aldrich, ref. 45511), an inhibitor of the ABA pathway.

All chemicals and a water control were freshly prepared to the desired concentrations in 0.5% of surfactant (“Calanque", Action Pin, Castets, France) to ensure the penetration of active molecules through the bud scales. Chemicals were sprayed on buds to runoff under a fume-hood and branches were left several minutes to allow them to dry. Branches were then transferred in the growth chamber (25°C, 16h light/ 8h dark, 60-70% humidity) in pots containing water. Bud break measurements were performed on flower buds as mentioned above.

### Phytohormones extraction

For each sample (see Table **S1**; Supplementary file at *Tree Physiology* online), 10 mg of frozen pulverised flower buds were weighed in a 2 mL tube. The extraction was carried out as previously described (Ali et al. 2018, Haddad et al. 2018, Lakkis et al. 2019) by adding 1 mL of cold 70% MeOH/ 29% H_2_O/1.0% formic acid, containing isotopically labelled internal standards. Then, the tubes were stirred at room temperature for 30 min and centrifuged (5427R, Eppendorf) at 16,000 rpm for 20 minutes at 4°C. The supernatant of each tubes were transferred into new tubes and evaporated to dryness using a Turbovap LV system (Biotage, Sweden). The dried extracts were dissolved with 1 mL of a 2% formic acid solution. The resuspended extracts were purified using a solid phase extraction (SPE) Evolute express ABN 1ml-30 mg (Biotage, UK). The eluate was evaporated to dryness and resuspended in 200 μL of 0.1% formic acid before analysis.

### Phytohormones quantification

ABA and conjugates (ABA-GE, PA, DPA) and GAs (GA_1_, GA_3_, GA_4_, GA_7_) were quantified by UHPLC-MS/MS as previously described (Lakkis et al. 2019). ABA, ABA-GE, gibberellins (GA_4_, GA_7_), [^2^H_6_]-ABA, and [^2^H_2_]-GA_4_ were purchased from OlchemIn (Olomouc, Czech Republic). DPA, PA, [^2^H_3_]-dihydrophaseic acid (D-DPA), and [^2^H_3_]-phaseic acid (D-PA) were purchased from National Research Council Canada (NRC, Saskatoon, Canada). Phytohormones were analyzed by an UHPLC-MS/MS system. The separation and detection were achieved using a Nexera X2 UHPLC system (Shimadzu, Japan) coupled to a QTrap 6500+ mass spectrometer (Sciex, Canada) equipped with an electrospray (ESI) source. Phytohormones separation was carried out by injecting 2 μL into a Kinetex Evo C18 core-shell column (100 × 2.1mm, 2.6μm, Phenomenex, USA) at a flow rate of 0.7 mL/min, and the column oven was maintained at 40°C. The mobile phases were composed of solvent A Milli-Q water (18 MΩ, Millipore, USA) containing 0.1% formic acid (LCMS grade, Fluka analytics, Germany), and solvent B acetonitrile LCMS grade (Fisher Optima, UK) containing 0.1% formic acid. The gradient elution started with 1% B, 0.0-5.0 min 60% B, 5.0-5.5 min 100% B, 5.5-7.0 min 100 % B, 7.0-7.5 min 1% B, and 7.5-9.5 min 1% B. The ionization voltage was set to 5kV for positive mode and -4.5 kV for negative mode producing mainly [M+H]^+^ and [M-H]^-^ respectively. The analysis was performed in scheduled multiple reaction monitoring (MRM) mode in positive and negative mode simultaneously with a polarity switching of 5 ms. All quantitative data were processed using MultiQuant software V 3.0.2 (Sciex, Canada). GA_1_, GA_3_ were not detected in the samples.

### Candidate gene identification

In order to select and investigate genes involved in the hormonal signaling pathways, we used the functional annotation tools Mercator4 (Schwacke et al. 2019) and eggnog (Huerta-cepas et al. 2017, 2019) on the sweet cherry ‘Regina’ genome (v1.0; Le Dantec et al. 2020). Results for the annotated sweet cherry genes were crossed-checked using pairwise sequence comparison with *Arabidopsis thaliana* proteins. Genes were identified in the obtained annotation database by key words and gene names from the literature, including Arabidopsis genes identified by Howe and colleagues (Howe et al. 2015). Details on the identified genes are available in Table **S2** (Supplementary file at *Tree Physiology* online).

### RNA-seq and differential expression analysis

Total RNA was extracted and sequenced as described in Vimont et al. (2019). Sequencing data are available online (Gene Expression Omnibus GSE130426). The quality of raw reads was assessed using FastQC (www.bioinformatics.babraham.ac.uk/projects/fastqc/) and possible adaptor contaminations and low quality trailing sequences were removed using Trimmomatic (Bolger et al. 2014). Raw reads sequences were previously mapped on the peach (*Prunus persica*) reference genome in a published analysis (Vimont et al. 2019). Here, we decided to remap the reads to the to the sweet cherry ‘Regina’ genome (v1.0; Le Dantec et al. 2020) using STAR (Dobin et al. 2013). Raw counts for each transcript were calculated using HTSeq (Anders et al. 2015). For the number of input reads and the percentage of mapped reads in each sample, please refer to Table **S1**. For each gene, Transcripts Per Million reads (TPM) were calculated (Wagner et al. 2012). TPM for the genes analysed in this study are available in the supplementary data file at *Tree Physiology* online. Differentially expressed genes (DEGs) for each combination of dormancy stages (pre-dormancy, endodormancy, dormancy breaking and ecodormancy) were assessed using DEseq2 R Bioconductor package (Love et al. 2014), in the statistical software R (R Core Team 2018), on filtered data. Genes with an adjusted *p-value* (padj) < 0.05 (Benjamini-Hochberg multiple testing correction method) and a log2Fold change > 1, in at least one of the comparisons, were assigned as DEGs (Table **S3**, Supplementary file at *Tree Physiology* online). We confirmed that the reads mapping from the different cultivars to the sweet cherry sequences of the candidate genes was similar in coverage (Fig. **S2**, Fig. **S3**), therefore supporting the robustness of the expression comparison between cultivars, expressed in TPM.

### Modelling

In order to explore the differences in the expression of ABA in the two cultivars, ‘Cristobalina’ and ‘Regina’, we took a mathematical modelling approach. We constructed a model incorporating information from the genes involved in the ABA signalling pathway.

Since NCEDs and CYP707As and UGT71B6 have been implicated in the production and catabolism of ABA, respectively, they were considered in the production and decay rates of ABA. ABA level at different times, t, for each cultivar is described by an ordinary differential equation:

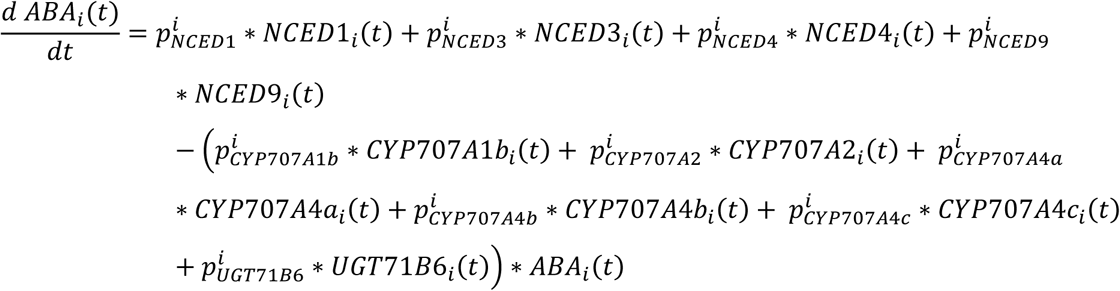

for i=1,2, where i=1 represents the index of cultivar ‘Regina’ and i=2 is the index of cultivar ‘Cristobalina‘.

In both cultivars, for the sake of simplicity, it was assumed that the rates are linearly dependent on the gene levels. For example, the rate of NCED1–dependent ABA production in Regina at a time t is described by 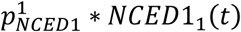 where 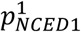 is a non-negative rate constant (model parameter). Genes (*NCED*s, *CYP707As* and *UGT*) are treated as the time-dependent parameters of the model and their values are taken from the data. More precisely, going back to the earlier example, the level of *NCED1* in ‘Regina’, labelled *NCED1*_1_(*t*), is a function of time t with values calculated from linearly interpolated mean ‘Regina’ data values of *NCED1*. Initial level of ABA at time 0 in each cultivar, i.e. ABA(0), is taken to be the mean level of ABA on the first day of measurement.

In order to show whether the differential in ABA in the two cultivars could be explained solely by the differences in *NCEDs, CYP7074As* and *UGT*, we tested whether there exists a set of parameters where the parameter values for both cultivar models are the same (i.e. 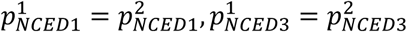 and so on) but for which the model simulations can show the ABA differences seen between the two cultivars. Latin Hypercube Sampling was used to select 100,000 parameter sets of X parameters (X being the number of genes used in the model). Since the gene levels can peak with level (in TPM) of up to 100 times higher than ABA levels (in ng/mg), the production and decay rate constants were bounded above by 0.001 and 0.005, respectively. We selected the parameter set with the lowest sum of the least squares as the best model for simulation and prediction. Once the model solutions were calculated, predictions for the best model were evaluated by comparing predicted ABA levels to measured ABA content in all ‘Regina’ and ‘Cristobalina’ samples using root mean square error (RMSE).

To explore which genes may have more or less significant effect on ABA levels during dormancy, we tested different combinations of genes, decreasing the number of genes from 10 to 3. Since testing all possible combinations would be very computationally expensive (for example, just selecting 7 out of 10 genes would require testing 10!/(7!×3!)=120 combinations of models), we decided to take a different approach. For the 9 gene models, we tested all nine model combinations where one gene is removed, and we selected the one with the lowest sum of the least squares as the best model for simulation and prediction, namely the 9 gene model that omits *PavCYP707A2* (Table **S4**, Supplementary file at *Tree Physiology* online). At the next step, when testing the 8 gene models, we removed *PavCYP707A2* and one of the other genes in turn. We tested now eight model combinations, repeating our steps above, down to the 3 gene models (see Table **S4** for more details on the tested combinations). Comparison of the best fitting models for each of the 3 to 10 gene models (with details listed in Table **S4**), revealed that a model with 6 genes had the lowest sum of the least squares overall. Finally, using the mean data measurements for *PavNCEDs, PavCYP7074As* and *PavUGT71B6* of the cultivar ‘Garnet’, we used this 6 gene model to predict the levels of ABA in the ‘Garnet’ cultivar. Since the initial value of ABA content in ‘Garnet’ cultivar was not measured, we took it arbitrarily to be 1 at the time 0 (this being a value that also falls within the range of the initial ABA levels of the ‘Regina’ and ‘Cristobalina’ cultivars).

## RESULTS

### Definition of the flower bud dormancy status of three cultivars with different flowering dates

We selected three sweet cherry cultivars based on their different dates of flowering: very early, early and late for ‘Cristobalina’, ‘Garnet’ and ‘Regina’, respectively. The stages of bud dormancy were defined based on anatomical and physiological elements (Fig. **2a**). We associated predormancy with developmental stages of the flower buds along green leaves and active growth (July to September). Dormancy onset is often hypothesized to occur at leaf fall (Chmielewski et al. 2017) and we observed leaf senescence at the beginning of October and complete leaf fall at the end of October. Therefore, dormancy onset was set in October and the beginning of endodormancy was established in November (Fig. **2a**). These stages were similar for all three cultivars. By definition, endodormancy correspond to the bud inability to fully develop under growth-inducing conditions while dormancy is considered as released when bud break is triggered by warm temperatures and/or long photoperiod (Lang et al. 1987). Consequently, dormancy status was assessed by forcing tests on branches carrying flower buds (Fig. **2b**). Endodormancy was defined for the dates with no bud break under forcing conditions (Fig. 2). Incomplete bud break within the population of flower buds indicated dormancy release stages while ecodormancy was defined by optimal bud break response to growth conditions (90-100% bud break; Fig. **2**). Here, the three cultivars were much contrasted in the timing of their dormancy phases, ‘Cristobalina’ exhibiting dormancy release on December 9^th^ (corresponding to a chill accumulation of 287 CH; Fig. **2c**), seven weeks earlier than ‘Garnet’ (January 29^th^; 628 CH; Fig. **2c**) and ten weeks earlier than ‘Regina’ (February 26^th^; 851 CH; Fig. **2c**).

**Figure 1.**
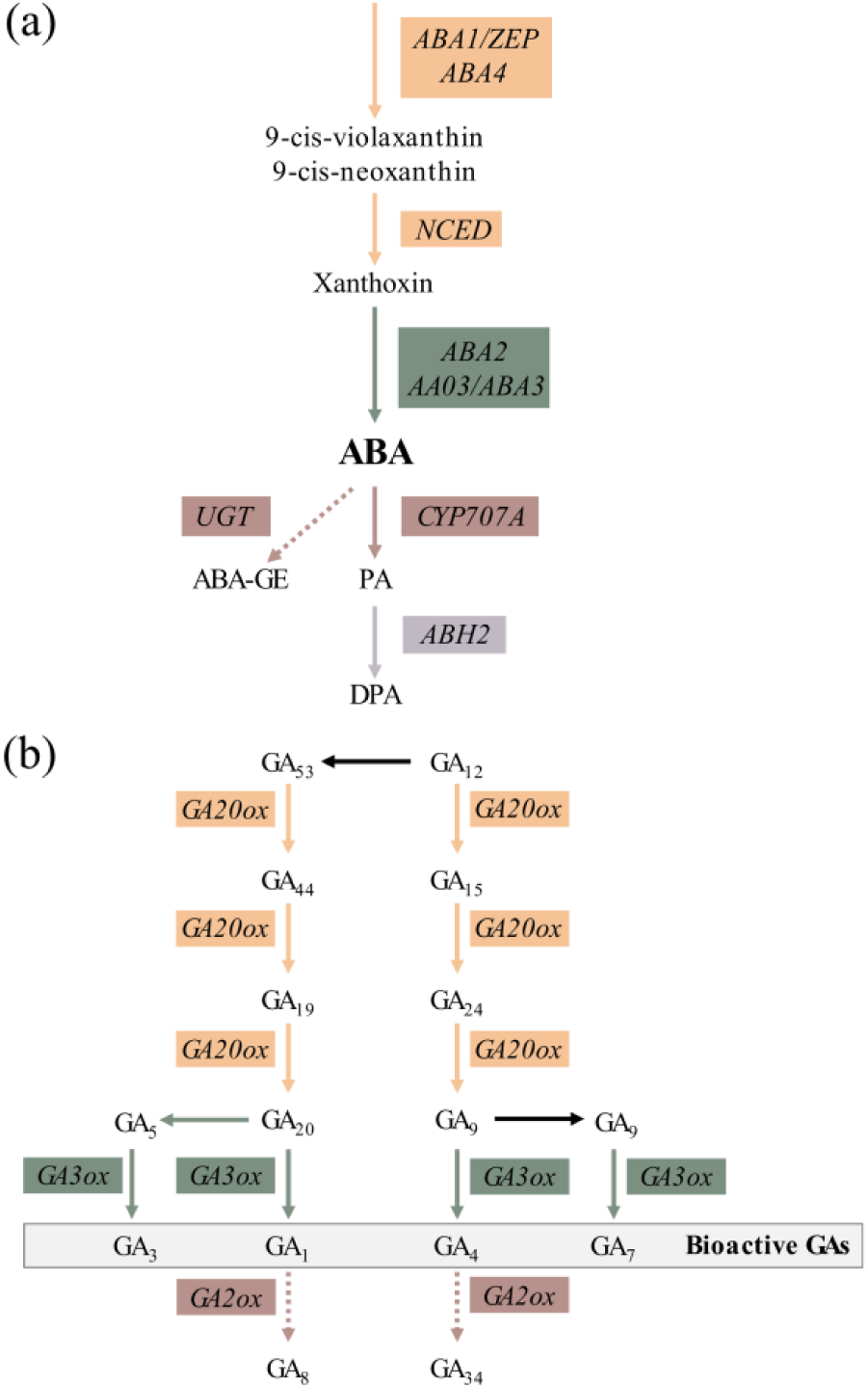
Biosynthesis and catabolism pathway for ABA and GAs. (**a**) ABA is synthesized through the action of five enzymes: zeaxanthin epoxidase (ZEP/ABA1), ABA-deficient4 (ABA4,) 9-cis epoxycarotenoid dioxygenase (NCED), alcohol dehydrogenase (ABA2) and short-chain dehydrogenase/reductase (AAO3/ABA3). ABA is mainly inactivated by ABA 8′-hydroxylase-catalyzed conversion to 8‘hydroxy ABA by cytochrome P450 monooxygenases, encoded by *CYP707A* (Nambara and Marion-Poll, 2005). 8‘hydroxy ABA is then spontaneously converted to phaseic acid (PA), which is further catabolized to dihydrophaseic acid DPA by a PA reductase (PAR) encoded by *ABA HYPERSENSITIVE2* (*ABH2*). ABA can be conjugated with glucose to inactive ABA-glucose ester (ABA-GE) by UDP-glycosyltransferases (UGT) (Dietz et al., 2000). (**b**) Bioactive GAs (GA_1_, GA_3_, GA_4_ and GA_7_) are synthetized by GA 20-oxidases (GA20ox) and GA 3-oxidases (GA3ox) and catabolized by GA 2-oxidases (GA2ox) (Yamaguchi, 2008). ABA: Abscisic acid; GA: Gibberellic acid.

**Figure 2.**
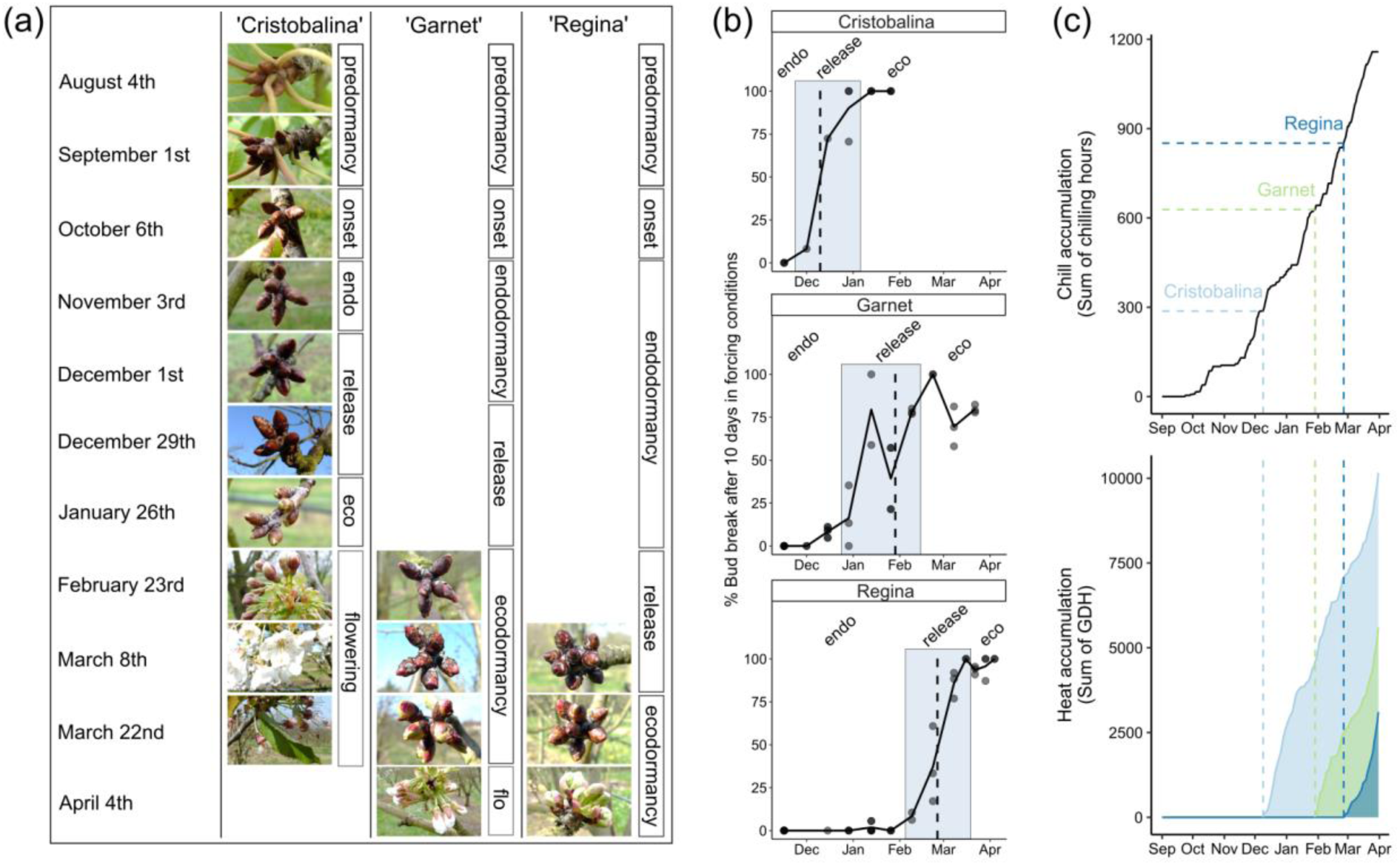
Dormancy status defined for the three sweet cherry cultivars. (a) bud stages at the sampling dates and the corresponding dormancy status as defined in the current study; (b) Evaluation of bud break percentage under forcing conditions was carried out for three sweet cherry cultivars displaying very early, early and late flowering dates: ‘Cristobalina’, ‘Garnet’ and ‘Regina’ respectively. The dotted line corresponds to the dormancy release dates, estimated at 50% of bud break after ten days under forcing conditions. Dots indicate the data points for the biological replicates; (c) Chill and heat accumulation for the three cultivars estimated using the dormancy release date and temperatures. Chill accumulation (calculated as the sum of chilling hours) stops after dormancy release, then followed by heat accumulation (calculated as the sum of growing degree hours, GDH).

### Exogenous GA application accelerates bud dormancy release

GA and ABA effect on the bud break response to forcing conditions was evaluated on branches carrying dormant flower buds as assessed by forcing tests on the late flowering cultivar ‘Fertard’ (Fig. **S1**). Results revealed that GA_3_ and GA_4_ both had significant dormancy alleviating effects, characterized by higher bud break percentage, which was confirmed for GA_3_ in a second experiment in 2018 (Fig. **3**, Fig. **S4**). However, no antagonist effect, namely bud break inhibition, was observed after a treatment with paclobutrazol, suggesting that GA biosynthesis was not required for the observed response to warm conditions.

In both seed and bud dormancy, it is hypothesized that GAs and ABA act antagonistically to control growth resumption and inhibition, respectively. We therefore tested the potential inhibiting effect of ABA on flower bud emergence. We did not observe a significant effect of ABA treatment on dormancy release, but bud break for ABA-treated branches was slightly higher than the control (Fig. **3**, Fig. **S4**). However, inhibiting ABA biosynthesis with fluridone activated bud break, consistent with the established role of ABA in promoting dormancy.

**Figure 3.**
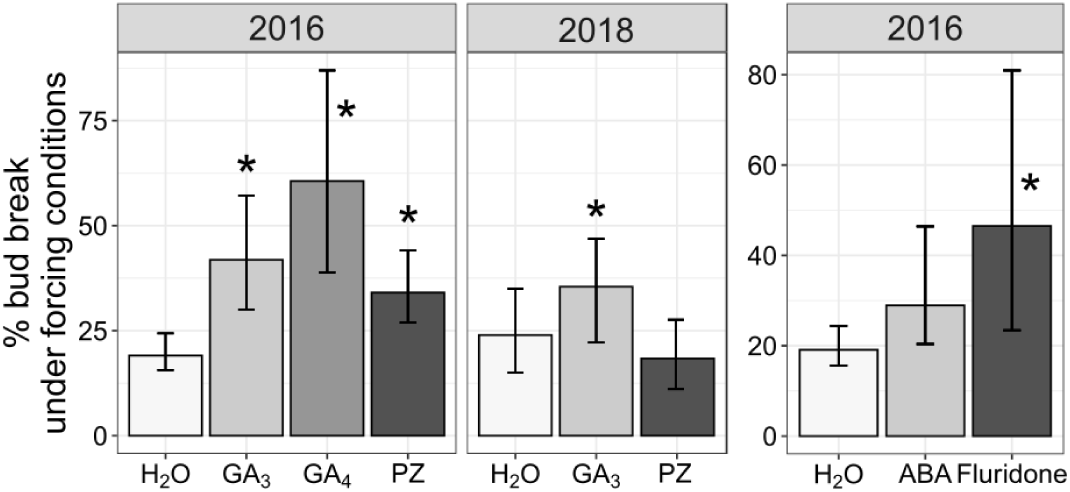
Effect of different GAs, ABA and their inhibitor on the sweet cherry dormancy status. Sweet cherry branches were treated with (a) 5 μM GA_3_, 5 μM GA_4_, 300 μM, paclobutrazol (GA pathway inhibitor), (b) 400 μM ABA and 5 μM fluridone (ABA pathway inhibitor) and transferred under forcing conditions (25°C, 60-70% humidity, 16 hours light). The percentage of flower bud break was recorded after 20 days. Error bars indicate the data range between the five biological replicates. Asterisks indicate treatments that differ significantly from untreated branches (Kruskal-Wallis test, p < 0.05). ABA: Abscisic acid; GA: Gibberellic acid.

### GA content changes during flower bud dormancy progression

In recent studies, distinct functions were identified for gibberellins during bud dormancy (Zhuang et al. 2013, Zheng et al. 2018). To test whether these results could be confirmed in sweet cherry buds, GA levels were determined over the whole bud development cycle in the very early and late flowering cultivars, ‘Cristobalina’ and ‘Regina‘. In details, we studied the content of bioactive GA_1_, GA_3_ GA_4_ and GA_7_ but the levels of GA_1_ and GA_3_ were undetectable in the samples. GA_4_, GA_7_ have a similar pattern over dormancy progression (Fig. **4**) rising from July onwards, which corresponds to flower primordia initiation and organogenesis. Bioactive GA levels also increase at the beginning of endodormancy (November – December), reaching their highest concentration during dormancy release. Results show a sharp decrease in the levels of GA_4_ and GA_7_ overlapping with ecodormancy for ‘Cristobalina’, which could not be observed for the late cultivar ‘Regina’, potentially due to the lack of ecodormancy coverage. Interestingly, the levels of GA_4_, and GA_7_ significantly differed between the two cultivars, especially during endodormancy and when dormancy release was triggered in the early cultivar (Fig. **S5b**). Notable differences in GA_7_ content were observed between these two cultivars during the entire time course, in which the late cultivar buds contained more GA_7_ than the early cultivar (Fig. **4**) while levels for GA_4_ were significantly higher in the early cultivar during endodormancy (Fig. **S5b**). Among the quantified active GAs, GA_4_ was detected at levels between three and eight times higher than GA_7_. GA_4_ was detected at higher levels in the early cultivar ‘Cristobalina’, with a relatively high level reached just after dormancy release in December. By contrast, levels of GA_7_ were higher in ‘Regina’, but with the maximal concentration measured just after dormancy release.

**Figure 4.**
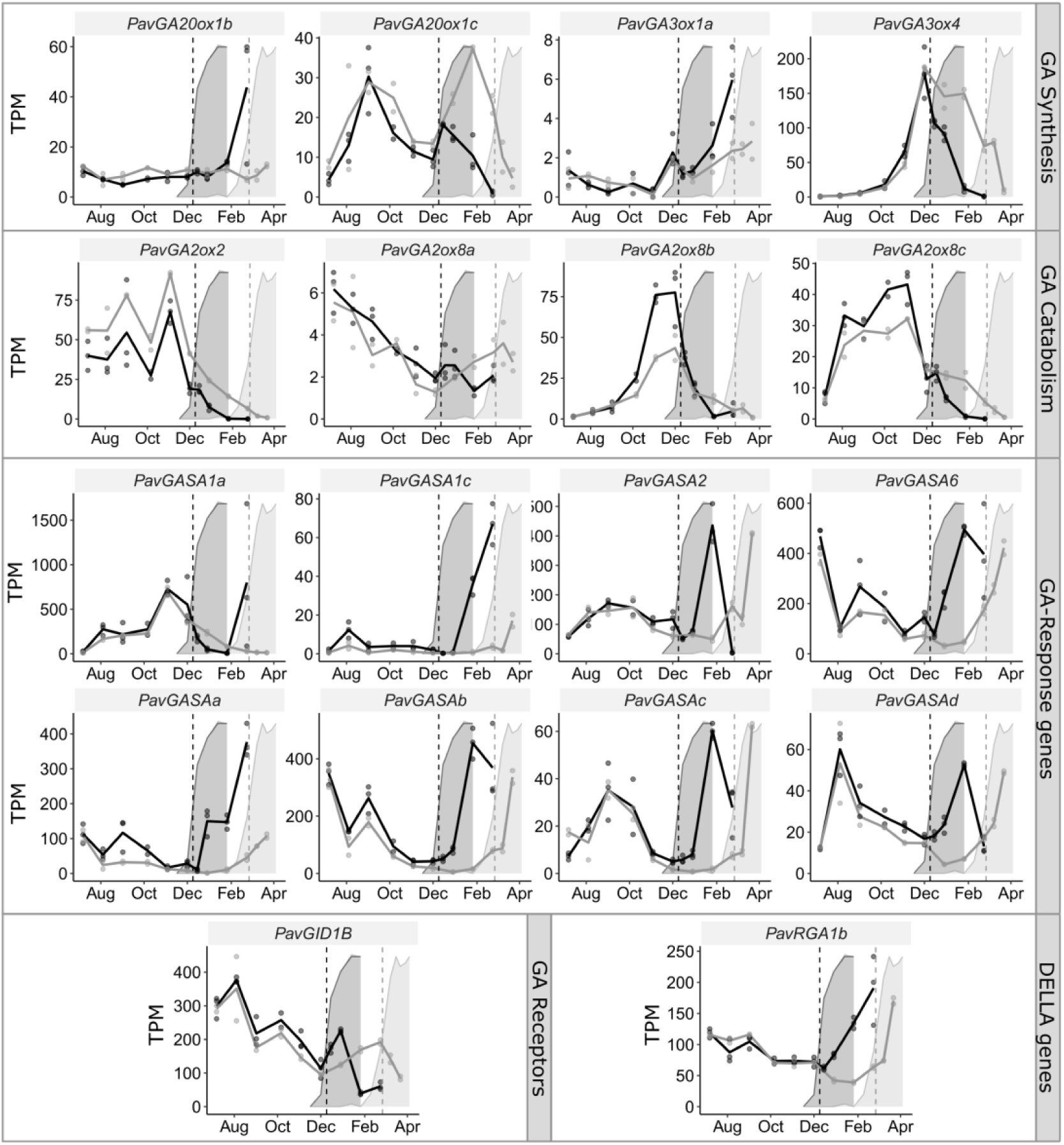
Levels of endogenous bioactive GA_4_ and GA_7_ in the flower buds of two sweet cherry cultivars during bud development. Black: ‘Cristobalina’, grey: ‘Regina‘. Background areas correspond to the dormancy depth evaluated as the percentage of bud break under forcing conditions (see Fig. 2b). Dotted lines represent dormancy release. GA: Gibberellic acid; Pre: Predormancy; O: dormancy onset; En: Endodormancy; R: Dormancy release; Ec: Ecodormancy.

### Expression of GA pathway-related genes have distinct patterns during sweet cherry bud dormancy

To better understand the mechanisms linked to the GA pathway during dormancy progression, we investigated genes involved in GA biosynthesis pathway, degradation, signal transduction and response. We found eleven *PavGA20ox* and five *PavGA3ox* for GA biosynthesis genes in the peach database (Table **S2**) but only four genes were differentially expressed over the dormancy period: *PavGA20ox1b, PavGA20ox1c, PavGA3ox1a* and *PavGA3ox4* (Fig. **5**, Table **S3**). Interestingly, *PavGA20ox1c* expression increased in December for both cultivars, regardless of their dormancy status but decreased prior to dormancy release. The marked increase in *PavGA20ox1c* expression could be correlated with the production of GAs around December (Fig. **4**). The last step of active GA biosynthesis relies on the activity of GA3ox essentially for the production of GA_1_ and GA_4_. Expression for *PavGA3ox4* increases as early as dormancy onset (October), followed by a sharp increase during endodormancy, reaching its highest expression value at maximum dormancy depth in December, for both cultivars (Fig. **5**). *PavGA3ox4* is then downregulated during or after dormancy release, with a marked lag between the two cultivars, potentially linked to their separate dormancy release date (Fig. **5**). Interestingly, *PavGA20ox1b* and *PavGA3ox1a* were both expressed during late ecodormancy, but with no obvious correlation with high GA levels (Fig. **4**). These results suggest that GA biosynthesis may not be activated after dormancy release, which is consistent with the previous observation that GA biosynthesis may not be needed for the initiation of ecodormancy. We identified four differentially expressed *PavGA2ox* genes, involved in GA inactivation (Fig. **5**, Tables **S2, S3**). *PavGA2ox2, PavGA2ox8a* and *PavGA2ox8c* genes were expressed before dormancy and during the early stages of dormancy. *PavGA2ox8b* and *PavGA2ox8c* were highly expressed during endodormancy, concomitantly with *PavGA3ox4* expression, thus suggesting a balance between synthesis and degradation that closely controls the levels of bioactive GAs.

In terms of GA signaling, our results show that the identified GA receptor-related *GA INSENSITIVE DWARF1B* (*PavGID1B*) is highly expressed during the predormancy (July, August) and early stages of dormancy (September, October; Fig. **5**). For ‘Cristobalina’, expression of the receptor gene sharply decreased after endodormancy was released. Ten GA-response genes, *GA Stimulated Arabidopsis* (*GASA*), potentially regulated by GAs (Aubert et al. 1998), were identified in the transcript dataset (Table **S2**) and we analyzed the eight genes differentially expressed during flower bud cycle (Fig. **5**, Table **S3**). Expression patterns are diverse but for the majority (*PavGASA1c, 6, a, b, c*, d), a decrease in expression was detected during deep dormancy (November, December), thus suggesting that GA-activated pathways are inhibited during dormancy, despite high contents in GAs (Fig. **4**). More strikingly, all *PavGASA* genes were sharply upregulated during dormancy release. However, one notable exception is *PavGASA1a* that is highly activated specifically during dormancy (Fig. **5**). The repression of GA by DELLA proteins is well characterized in annuals (Zentella et al. 2007), so to further investigate GA pathway, we identified five genes coding for predicted DELLAs, namely *PavRGA1a, b, PavRGL3a, b, c* (Table **S2**). Among them, only *PavRGA1b* was differentially expressed during bud development, characterized by a marked expression increase during dormancy release and ecodormancy (Fig. **5**).

**Figure 5.**
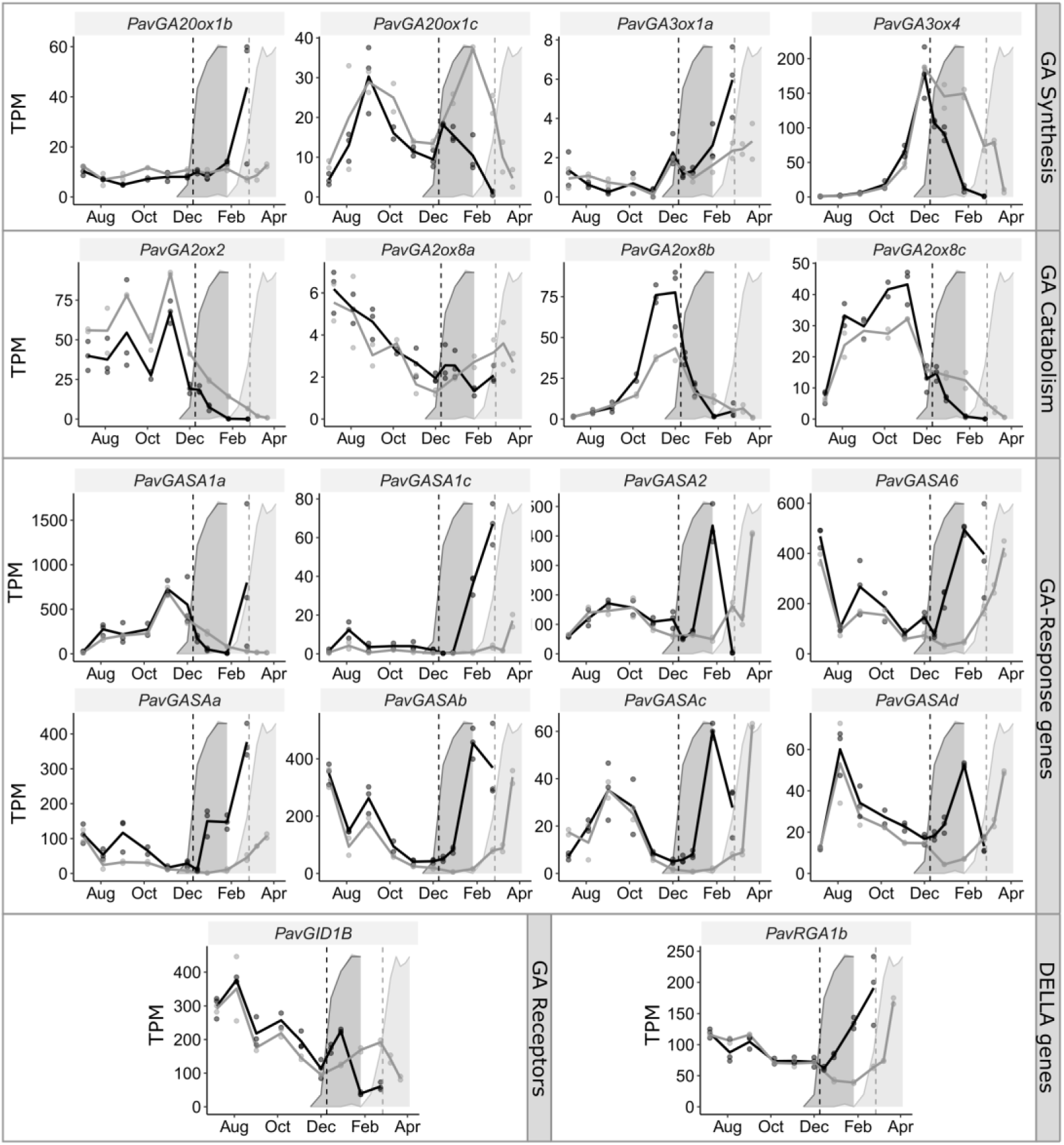
Transcriptional dynamics of genes associated with GA pathway in the flower buds of two sweet cherry cultivars during bud development. Expression of specific genes involved in GA biosynthesis pathway, degradation, signal transduction and response are represented in TPM (Transcripts Per Million reads). Black: ‘Cristobalina’, grey: ‘Regina‘. Background areas correspond to the dormancy depth evaluated as the percentage of bud break under forcing conditions (see Fig. 2b). Dotted lines represent dormancy release.GA: Gibberellic acid; GA20ox: GA 20-oxidases, GA3ox: GA 3-oxidases; GA2ox: GA 2-oxidases; GID: GA INSENSITIVE DWARF; GASA: GA Stimulated Arabidopsis; RGA: REPRESSOR OF GA.

**Figure 6.**
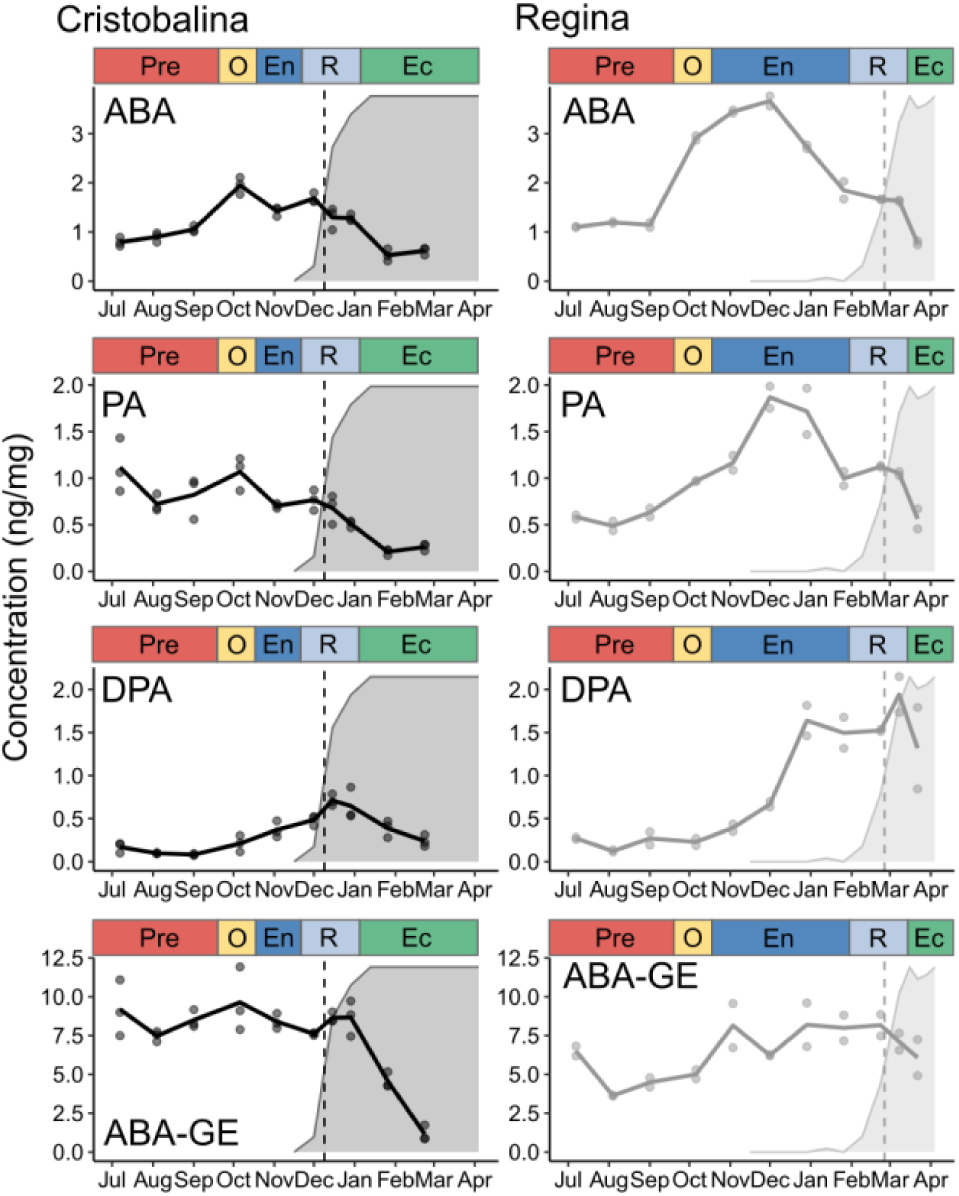
Levels of endogenous bioactive ABA and ABA conjugates in the flower buds of two sweet cherry cultivars during bud development. Black: ‘Cristobalina’, grey: ‘Regina‘. Background areas correspond to the dormancy depth evaluated as the percentage of bud break under forcing conditions (see Fig. 2b). Dotted lines represent dormancy release. ABA: Abscisic acid; PA: phaseic acid; DPA: dihydrophaseic acid, ABA-GE: ABA-glucose ester; Pre: Predormancy; O: dormancy onset; En: Endodormancy; R: Dormancy release; Ec: Ecodormancy.

### ABA levels rise at the onset of dormancy

Several studies have highlighted a strong correlation between ABA content and dormancy status and to address this issue in sweet cherry flower buds, we measured ABA levels, as well as PA and DPA, which are catabolites of ABA, in both cultivars. Results show an increase in ABA and PA content during the early stages of dormancy, reaching their highest levels in October and December for ‘Cristobalina’ and ‘Regina’, respectively, which is approximately two months prior to dormancy release for both cultivars. This ABA peak is followed by a decrease in ABA levels, accompanied by increased levels of DPA, preceding dormancy release (Fig. **6**). ABA, PA and DPA levels detected during dormancy for the early cultivar ‘Cristobalina’ are significantly lower than for the late cultivar (Fig. **S5a**). We can therefore hypothesize that dormancy depth may be correlated with ABA contents in sweet cherry buds.

Esterification of ABA with glucose was also monitored and the concentrations of ABA-GE were higher in both cultivars compared with ABA and its conjugates. The ABA-GE content was constantly high in ‘Regina’ over the whole cycle, with a slight increase during endodormancy induction, while it markedly decreased in ‘Cristobalina’ during ecodormancy (Fig. **6**, Fig. **S5a**). However, this observation could be due to a low coverage of ecodormancy in the ‘Regina’ samples. Throughout pre-dormancy stages, ABA-GE content was significantly higher in the early cultivar ‘Cristobalina’ compared to the late cultivar ‘Regina’ (Fig. **S5a**).

**Figure 7.**
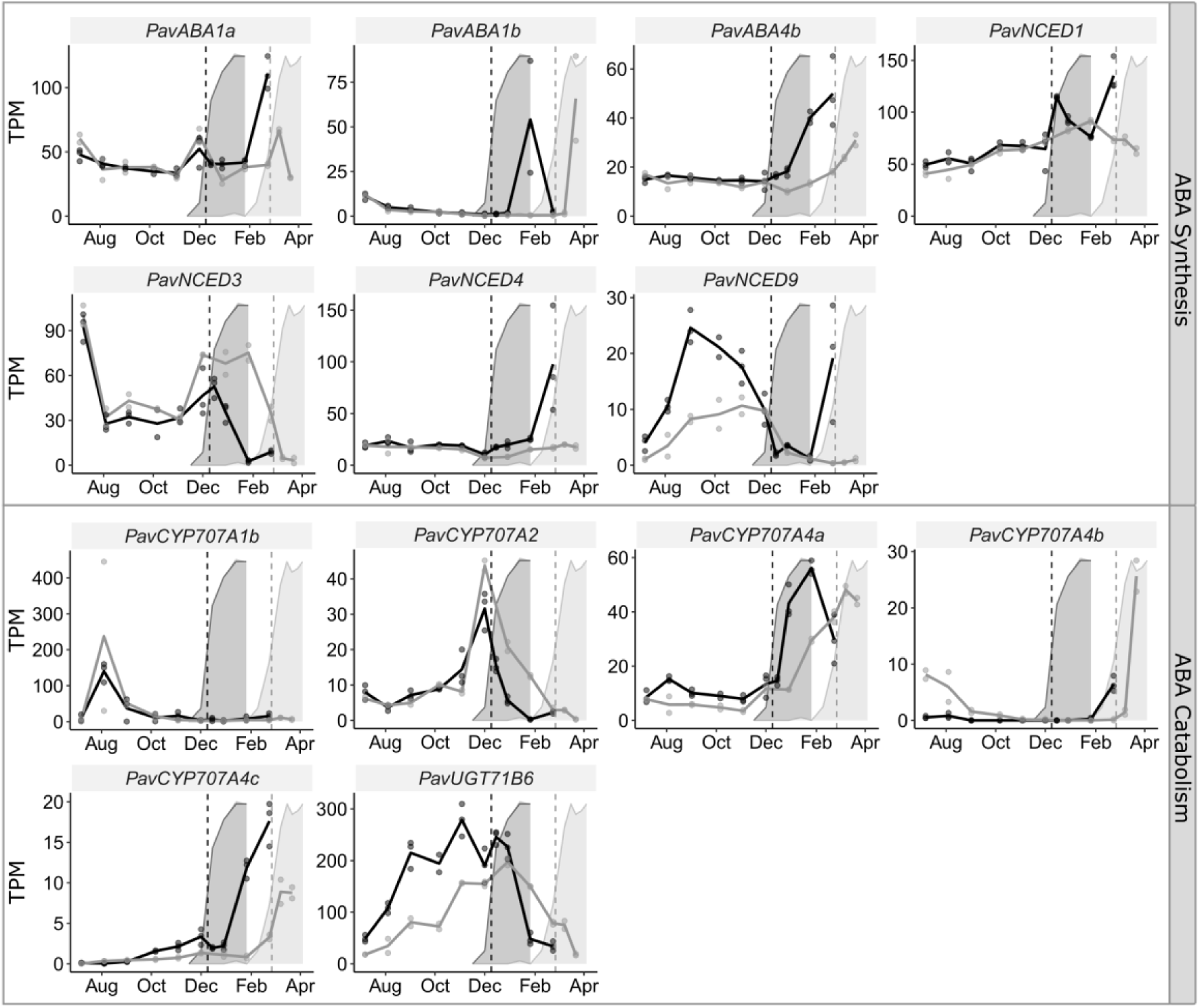
Transcriptional dynamics of genes associated with ABA biosynthesis and catabolism in the flower buds of two sweet cherry cultivars during bud development. Expression of specific genes are represented in TPM (Transcripts Per Million reads). Black: ‘Cristobalina’, grey: ‘Regina‘. Background areas correspond to the dormancy depth evaluated as the percentage of bud break under forcing conditions (see Fig. 2b). Dotted lines represent dormancy release. ABA: Abscisic acid; NCED: 9-cis epoxycarotenoid dioxygenase; UGT: UDP-GLYCOSYLTRANSFERASE.

### Analysis of genes involved in ABA pathway

We further explored the genes involved in ABA synthesis and catabolism. Expression for genes involved in the multiple ABA biosynthesis steps, *PavABA1a, PavABA1b, PavABA4b, PavNCED1* and *PavNCED4*, is not correlated with ABA levels while expression patterns for *PavNCED3* and *PavNCED9* genes seem strongly linked to ABA contents and dormancy status (Fig. **7**). In particular, *PavNCED9* expression shows a sharp increase during dormancy onset and a marked decay before dormancy release. On the other hand, low ABA levels are associated with increased expression for ABA catabolism genes *PavCYP707A1b* and *PavCYP707A4s* before and after dormancy respectively (Fig. **7**). By contrast, *PavCYP707A2* is characterized by a sharp increase in December, followed by a marked decrease during dormancy release in ‘Cristobalina’ but before induction of dormancy release in ‘Regina‘. In addition, we identified *PavUGT71B6* (Supporting Information Table **S2**), a sweet cherry ortholog of the *Arabidopsis thaliana UDP-glycosyltransferase 71B6*, that preferentially glycosylates ABA into ABA-GE (Priest et al., 2006). *PavUGT71B6* was considerably upregulated in the early cultivar compared to the late cultivar, with a gradual increase between July and deep dormancy followed by a decrease in expression during endodormancy release (Fig. **7**).

**Figure 8.**
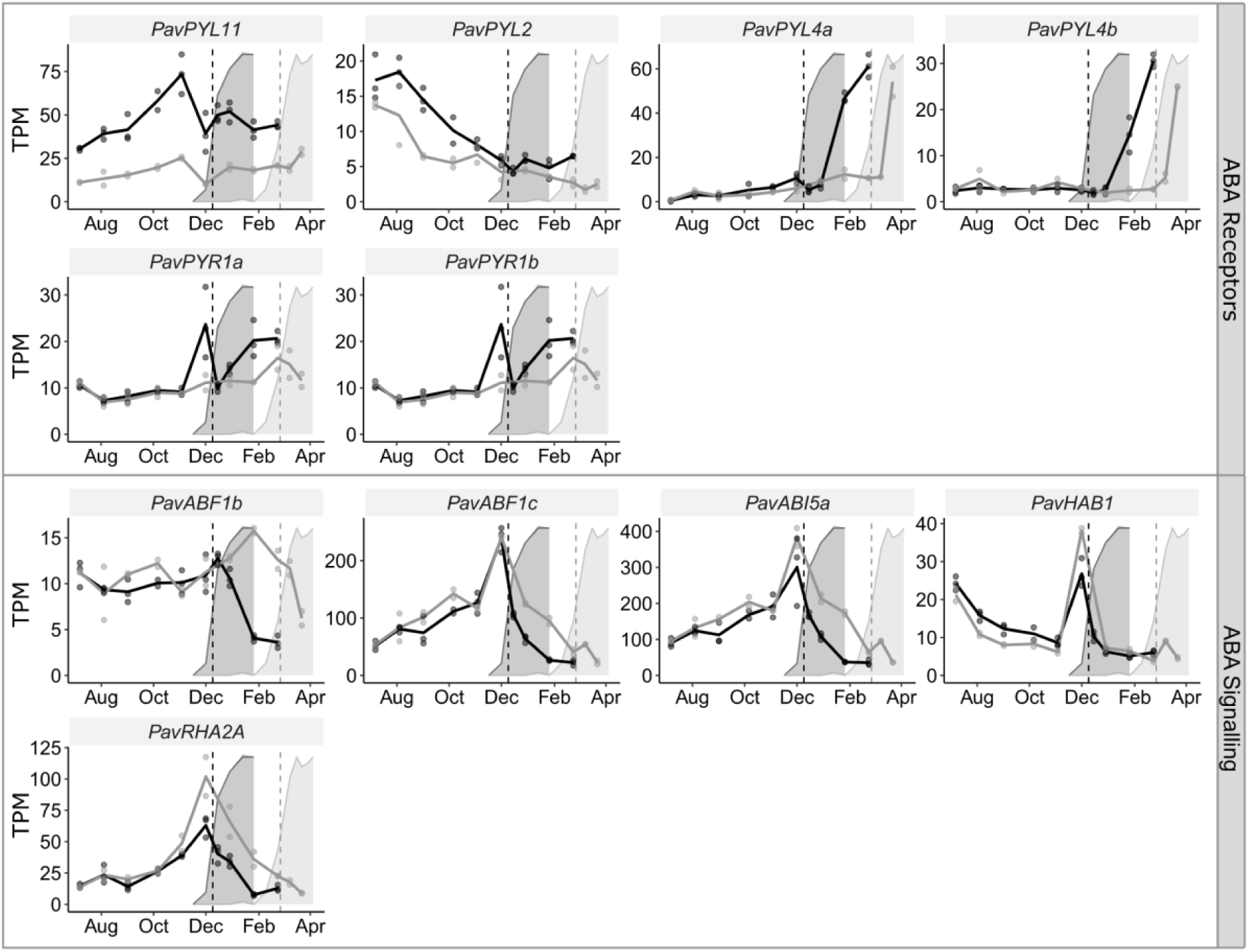
Transcriptional dynamics of genes associated with ABA signalling in the flower buds of two sweet cherry cultivars during bud development. Expression of specific genes are represented in TPM (Transcripts Per Million reads). Black: ‘Cristobalina’, grey: ‘Regina‘. Background areas correspond to the dormancy depth evaluated as the percentage of bud break under forcing conditions (see Fig. 2b). Dotted lines represent dormancy release. ABA: Abscisic acid; PYR: PYRABACTIN RESISTANCE; PYL: PYR-like; ABF: ABSCISIC ACID RESPONSIVE ELEMENTS-BINDING PROTEIN; ABI; ABA INSENSITIVE; HAB: homology to ABI2; RHA2A: RING-H2 A.

In addition, we examined sweet cherry gene predictions for genes involved in ABA signaling (Fig. **8**). Among the eight ABA receptors *PYR*/*PYL* genes identified for sweet cherry (Table **S2**), six were differentially expressed during flower bud dormancy (Fig. **8**, Table **S3**). In particular, *PavPYL11* expression was correlated with ABA levels in ‘Cristobalina’, increasing after dormancy onset and decaying after dormancy release (Fig. **8**), but not in ‘Regina‘. However, for the *PavPYL2* and *PavPYL4a/b* genes, the expression was activated before and after endodormancy, respectively. In terms of ABA-mediated signals, differentially expressed ABA response genes, *PavABF1c, PavABI5a, PavHAB1* and *PavRHA2A*, had similar patterns with a sharp increase in expression in December and a marked inhibition after the peak (Fig. **8**).

**Figure 9.**
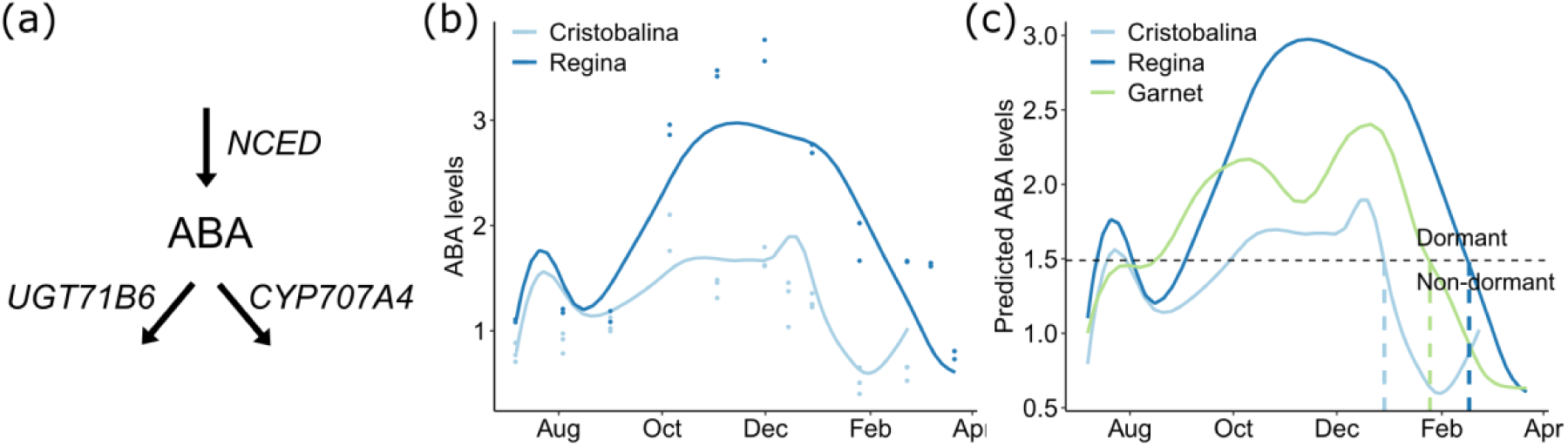
Modelling ABA. (**a**) Conceptual model used to simulate ABA content. ABA synthesis is controlled by PavNCED1, PavNCED3, PavNCED4 and PavNCED9 proteins; ABA is deactivated by 8‘-hydroxylases PavCYP707A4 and PavUGT71B6. The assumption is that enzymatic activity is proportional to gene expression levels. (**b**) Simulated content of ABA using the model (lines) with means of data (circles) for cultivars ‘Cristobalina’ and ‘Regina‘. The best model was obtained for simulation with parameters: 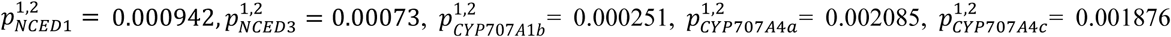 and 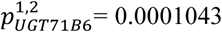 (**c**) Simulated levels of ABA for cultivars ‘Cristobalina’ and ‘Regina’, that were used to calibrate the model, and for ‘Garnet‘. Arbitrary level of ABA set at 1.49 ng/mg (black dash line) is reached by ‘Cristobalina’, ‘Garnet’ and ‘Regina’ simulations on December 29^th^, January 25^th^ and February 17^th^, respectively (colored dash lines). ABA: Abscisic acid; NCED: 9-cis epoxycarotenoid dioxygenase; UGT: UDP-GLYCOSYLTRANSFERASE.

### Modelling suggests ABA levels control onset and duration of dormancy

Based on observations that ABA levels are correlated with dormancy depth, ABA content has been proposed as an indicator to assess dormancy status in sweet cherry (Chmielewski et al. 2017). We further investigated the dynamics of ABA biosynthesis and catabolism to estimate dormancy onset and duration. First, we assumed that ABA synthesis is mainly controlled by *PavNCED1, 3, 4* and 9 while ABA is converted to PA by 8‘-hydroxylases *PavCYP707A* and esterified into ABA-GE by *PavUGT71B6* (Fig. **9a**). Since data for protein activity are not available, we used transcript levels as a proxy for enzymatic activity. We then used an Ordinary Differential Equation (ODE) approach to model how dormancy may be regulated by ABA levels. Among all tested gene combinations, the best model simulates ABA content based on the expression of only six genes, namely *PavNCED1, PavNCED3, PavCYP707A1b, PavCYP707A4a, PavCYP707A4c* and *PavUGT71B6* (Table **S4**, Fig. **S6**). The mathematical model, based on identical parameter set for both cultivars, shows a good fit to the data, with a global RMSE equal to 0.38 for all data, and 0.33 and 0.44 for ‘Cristobalina’ and ‘Regina’ respectively indicating that the differential in ABA levels between the two cultivars may be explained by the differences in gene expression of the relevant enzymes (Fig. **9b**). To validate the model, we simulated ABA levels for a third cultivar ‘Garnet’ that was examined along ‘Cristobalina’ and ‘Regina‘. Endodormancy was released on January 21^st^ for ‘Garnet’, an early flowering cultivar. Based on expression data for *PavNCED1* and *3, PavCYP707A1b, PavCYP707A4a, PavCYP707A4c* and *PavUGT71B6* genes (Fig. **9**, Fig. **S7**), the model simulated ABA levels for ‘Garnet’ during dormancy (Fig. **9c**). Simulated ABA content for ‘Garnet’ increases during dormancy onset, reaches high values during endodormancy and decreases before dormancy release, later than for the very early cultivar ‘Cristobalina’, and earlier than for the late cultivar ‘Regina‘. Highest estimated levels for ABA are lower in ‘Garnet’ than in ‘Regina’ but higher than ‘Cristobalina‘.

Observed and simulated levels of endogenous ABA at the date of dormancy release show a good match for both ‘Cristobalina’ and ‘Regina’ cultivars (Fig. **6**, Fig. **9b**). ABA levels are low before dormancy onset but as they increase dormancy is triggered; high ABA levels maintain dormancy but they decrease under chilling temperatures and endodormancy is released as ABA content falls. Model simulations show a peak in July, which is missing from the measured ABA levels. The Garnet ABA simulated levels lie between those of ‘Cristobalina’ and ‘Regina’ levels, thus if ABA content is related to dormancy release, then our modelling predicts that under a chosen fixed threshold of ABA, Garnet dormancy release will occur between the other two release dates. For example, ABA simulated levels of ‘Cristobalina’, ‘Regina’ and 'Garnet' would fall below the average threshold of 1.49 ng/mg on December 29^th^, February 17th and January 25^th^, respectively. Choice of any other threshold during this period will result in the same date order. Such ordering is observed in our data for dormancy release (Fig. **2a**); ‘Cristobalina’ and ‘Regina’ release dates are December 9^th^ and February 26^th^, while 'Garnet' release date is the January 29^st^; thus opening the possibility that there is a threshold for ABA concentration that determines the dormancy status. Since we can account for the dormancy behavior of different cultivars based on the expression of a small number of key genes regulating ABA levels, this underscores the central role of this phytohormone in the control of dormancy progression.

## DISCUSSION

### Sweet cherry specific GA signaling during bud dormancy

We have shown that GA_3_ and GA_4_ have significant dormancy alleviating effects in sweet cherry cultivars, similar to previous observations in hybrid aspen (*Populus tremula* × *Populus Tremuloides*; Rinne et al. 2011), Japanese apricot (*Prunus mume*; Zhuang et al. 2013) and peach (Donoho and Walker 1957), but opposed to null or inhibitory effects on dormancy release previously reported in hybrid aspen and grapevine (Rinne et al. 2011, Zheng, et al. 2018). In addition, high levels of GA_4_ and GA_7_ were detected in sweet cherry flower buds, similarly to results obtained in pear (Ito et al. 2019), whereas GA_1_ and GA_3_ were undetectable in the samples, contradictory to the high GA_1_ and GA_3_ levels recorded in grapevine and Japanese apricot buds during dormancy (Zhang et al. 2018, Zheng et al. 2018). These observations are consistent with the hypothesis that GAs act in a complex manner, with differential effects depending on concentrations and developmental phases, as discussed in Zheng et al. (Zheng et al. 2018). In addition, GA effects might be species or even cultivar-specific, with complex interactions with environmental factors, such as temperature or photoperiod. Further investigation on the regulation of the GA-related pathway during dormancy in sweet cherry, under contrasted environmental conditions, will bring key elements to the current hypotheses. Furthermore, given the complexity of GA quantification, it is possible that key GA dynamics might have been missed in the present study and therefore, expression of GA-pathway-related genes might be more reliable. Here, based on our results, we propose our hypotheses on GA signaling involving different transcriptional cascades during sweet cherry flower bud dormancy:

i. During predormancy stages (July to September), GA_4_ and GA_7_ content progressively increases, associated with an increase in *PavGA20ox1c* expression, and the activation of GA signaling, as revealed by the increased expression of GA-response genes *PavGASA2, 6, b, c* and *d*. Enhanced GA content potentially triggers the expression of *PavGA2ox8c* to catalyze GA deactivation as part of the feedback regulation (Yamaguchi 2008). In Arabidopsis, proteins GASA4 and GASA6 are involved in flower development and cell elongation, respectively, in response to GA signaling (Roxrud et al. 2007, Zhong et al. 2015) therefore we can hypothesize that gibberellin signaling, mostly driven by *PavGA20ox1c* in this phase may control flower bud organogenesis and development.
ii. After endodormancy is induced, there is a marked increase in GA_4_ and GA_7_ contents. Gibberellin homeostasis seems to be actively controlled during endodormancy through enhanced expression of biosynthesis gene *PavGA3ox4* and deactivation genes *PavGA2ox8b* and *PavGA2ox8c*. Consistently, the absence of paclobutrazol effect on bud break suggest that GAs are synthesized prior to dormancy release. Interestingly, previous studies have shown that cellular transport is blocked during dormancy (Rinne et al. 2011, Tylewicz et al. 2018) and we can therefore hypothesize that although GAs may be present in the bud, their growth-promoting effect is physically inhibited. In agreement with acute GA deactivation, only one GA-response gene is activated during this phase, *PavGASA1a*, thus suggesting a very specific response. Expression for the five GA-related genes up-regulated during endodormancy sharply decrease during dormancy release, thus supporting the hypothesis that this is an endodormancy-specific regulation.
iii. As observed in the early cultivar ‘Cristobalina’, levels for GA_4_ and GA_7_ decrease during ecodormancy, associated with low, if not null, expression for *PavGA20ox1c* and *PavGA3ox4* synthesis genes, as well as *PavGA2ox8b, PavGA2ox8c, PavGA2ox* deactivation genes and *PavGID1B* receptor. Interestingly, the GA-response *PavGASA* genes and the DELLA gene *PavRGA1b* are markedly activated during ecodormancy, thus suggesting that GA-stimulated pathways are up-regulated. Indeed, despite decreasing GA_4_ and GA_7_ levels during ecodormancy, GA-response pathways seem to be mostly activated when *GA2ox* genes are down-regulated after dormancy release so further investigation on the control of GA deactivation during dormancy could unravel a potential regulation by cold accumulation. In addition, bioactive GAs, including GA_1_ and GA_3_, that were not detected in the current study might be key actors in the growth resumption stage of ecodormancy in sweet cherry. Acute GA response occurring during ecodormancy, including activation of *GASA* genes, was previously reported in grapevine (Zheng et al. 2018), oak (Ueno et al. 2013), pear (Yang et al. 2019) and Japanese apricot (Zhang et al. 2018). Despite the fact that DELLA genes’ activity is mainly regulated by protein stability, the up-regulation of *PavRGA1b* is consistent with the GA response during ecodormancy, associated with a decrease in *PavGIDB* expression. These results suggest that the GA homeostasis, critical during active growth, is controlled in sweet cherry by the DELLA proteins, that target GA biosynthesis and receptor genes and impacts GA balance through a feedback regulation, as previously shown in Arabidopsis and rice (Gagne et al. 2002, Ueguchi-Tanaka et al. 2007, Zentella et al. 2007).

### Dormancy depth is correlated with endogenous ABA content

Exogenous application of ABA on the sweet cherry dormant bud showed that buds at this specific stage were not affected by ABA treatment, in contrast to the observation that ABA treatment had a significant effect on grapevine bud break (Zheng et al. 2015). However, as shown by the high expression of genes involved in ABA degradation, including *PavCYP707A2*, high catabolism ability during endodormancy might have limited the effect of exogenous ABA. It might also suggest that ABA response is saturated in the context of very high ABA levels during endodormancy, therefore limiting the effect of additional ABA. Nevertheless, we observed that dormancy release was triggered in buds treated with fluridone that inhibits ABA biosynthesis, as previously reported in pear (Yang et al. 2020), thus suggesting that high ABA levels may act to maintain dormancy. This was further confirmed by the observed elevated ABA levels correlated with endodormancy in both cultivars, as well as the differences in ABA levels between the early and late flowering cultivars. Our results are consistent with the hypothesis that dormancy is triggered and maintained when ABA levels are above a threshold, taking into account the potential heterogeneity within a population of flower buds. Bud dormancy may subsequently be released if ABA levels fall below the threshold. Consequently, such a dormancy release threshold would be reached earlier in the season for early cultivars with less ABA (Wen et al. 2016) or when ABA levels are lower due to various environmental conditions (Chmielewski et al. 2017). Recent reports have indeed shown how low or high levels of ABA closely drive the depth of dormancy by controlling the blockage of cellular transports (Tylewicz et al. 2018, Singh et al. 2019).

We show that during the predormancy and dormancy onset stages, ABA levels are correlated with an increase in the expression of *PavNCED9*, then *PavNCED3* genes in both cultivars. Consistently with elevated ABA levels, genes encoding ABA receptors (*PavPYL12, 8*), as well as ABA response genes, are highly expressed during deep dormancy. One interesting result is that *PavNCED9* expression peak coincides with the highest ABA levels in October in ‘Cristobalina’ and in December in ‘Regina’, approximately two months before dormancy is released. This suggests that the deepest dormancy state might occur earlier in ‘Cristobalina’ than in ‘Regina’ and therefore question whether endodormancy is indeed induced simultaneously in both cultivars. Further physiological observations during the early stages of dormancy induction, including flower primordia developmental context (Fadón et al. 2018), could help better understanding the observed differences. Afterwards, genes involved in ABA degradation (*CYP707A*) are highly expressed during dormancy release and positively correlated with a decrease of ABA content. Similar results were reported in other *Prunus* species (Zhang et al. 2015, Wang et al. 2016, Tuan et al. 2017), thus supporting the hypothesis that ABA signaling pathways play a major role in the regulation of dormancy induction, maintenance and release.

Presently, a remaining question to elucidate is what drives the decrease in ABA levels around dormancy release. Which mechanisms are involved in the down-regulation of *PavNCEDs* and up-regulation of *PavCYP707As* through chill accumulation? Firstly, several reports indicate that ABA might regulate its own accumulation and high levels of ABA attained during endodormancy could up-regulate the expression of catabolic genes such as *PavCYP707A4*, leading to a global decrease in ABA content. Secondly, *DORMANCY-ASSOCIATED MADS-BOX* (*DAM*) genes have been strong candidates for a key role in dormancy promotion and maintenance (Rodriguez et al. 1994, Bielenberg et al. 2008). *DAM* genes are highly expressed during dormancy and their expression is inhibited by chill accumulation (Jiménez et al. 2010, Hao et al. 2015), but more interestingly, recent studies have highlighted the direct effect of *DAM* on the activation of ABA biosynthesis (Tuan et al. 2017, Yamane et al. 2019). Further *in vitro* assays suggest a potential regulation between CBF proteins and *DAM* promoters (Niu et al. 2015, Zhao et al. 2018), which could suggest that when *CBF* genes are transiently activated in response to low temperatures, they modulate the expression of *DAM* genes that subsequently upregulate *NCED* genes and ABA biosynthesis in the first stages of dormancy. However, this may not explain the inhibiting effect of prolonged cold temperatures on *DAM* genes expression and ABA levels. Subsequently, more similarly to mechanisms controlling vernalization in annual plants (Horvath 2009), chill accumulation may induce chromatin modifications that silences *DAM* genes (Leida et al. 2012, Ríos et al. 2014, de la Fuente et al. 2015) and inhibits ABA production, consistently with decreasing ABA levels observed as soon as January. Interestingly, a recent study conducted in hybrid aspen shows that chilling and long photoperiods induces a decrease in ABA levels, which subsequently drives the decrease in *SHORT VEGETATIVE PHASE-LIKE* (*SVL*) expression (Singh et al. 2018, 2019). *SVL* shows similarity to *SHORT VEGETATIVE PHASE* (*SVP*) and *DAM* genes and this mechanism suggests that ABA might also act on the expression of *DAM* genes. Regulation of *DAM* and ABA pathways by long term cold temperatures should be further investigated to better understand how temperature variations control dormancy progression in sweet cherry flower buds.

### A contrasted ABA synthesis and conjugation balance between early and late cultivars

Overall, although the dynamics of expression for *PavNCED* and *PavCYP707A* genes effectively explain the increasing and decreasing pattern of ABA levels between dormancy induction and release, we further investigated whether they could account for the significant differences observed for ABA levels between the two cultivars. Contrasted expression patterns for *PavNCED3, PavCYP7074a* and *PavCYP7074c* were not sufficient to explain the noticeably contrasted ABA accumulation during endodormancy between the two cultivars. Differences in ABA catabolites between cultivars were also observed. While PA content followed the same pattern as ABA, DPA was particularly high during dormancy release, inversely correlated with the decreasing ABA levels. Interestingly, Weng and colleagues (Weng et al. 2016) exposed the compensatory effect of PA under low ABA conditions, in which PA is recognized by ABA receptors (PYL5 and 2) allowing a supplementary growth inhibition effect. In sweet cherry, increased *PavCYP707A2* expression may explain the higher levels of PA during dormancy in ‘Regina’ and might therefore result in even deeper dormancy by the inhibitory combination of ABA and PA. Since free ABA levels may be controlled through conjugation as well (El Kayal et al. 2011, Chmielewski et al. 2018, Liu and Sherif 2019), we investigated the conversion of ABA to ABA-GE and we found a *PavUGT71B6* gene characterized by strikingly higher expression levels for the early cultivar compared to the late cultivar. UGT71B6 orthologs in Arabidopsis and Adzuki bean act specifically for ABA conjugation into ABA-GE (Xu et al. 2002, Priest et al. 2006), so up-regulation of *PavUGT71B6* expression in ‘Cristobalina’ during dormancy may explain the higher content of ABA-GE. We can therefore hypothesize that the low ABA content in the early cultivar may be due to active catabolism of ABA to ABA-GE.

### Towards new phenology approaches based on molecular mechanisms

Following our observations that ABA levels were correlated with dormancy status and that dynamics of expression for ABA synthesis and catabolism may explain the differences observed between cultivars, we have successfully modeled ABA content and dormancy behavior in three cultivars exhibiting contrasted dormancy release dates. Indeed, ABA had been proposed as an indicator for dormancy release in sweet cherry (Chmielewski et al. 2017) but to our knowledge, this is the first attempt to simulate dormancy onset and duration using molecular data. Only a small number of key genes regulating ABA were sufficient to account for all variations in ABA levels and dormancy progression overtime and between cultivars. However, although we could verify the model on dormancy release dates in one independent cultivar, additional experiments with precise ABA levels evaluation will be necessary to validate the actual prediction of ABA levels based on gene expression associated with the dormancy status in ‘Garnet‘. In addition, previous analyses have shown that ABA levels are highly variables between years (Chmielewski et al. 2017) therefore further analyses are needed to explore and validate the current model.

Antagonistic actions of ABA and GA have been extensively studied in seed dormancy (Shu et al. 2018) and the ABA/GA ratio is often proposed as a determinant factor in the control of rest and growth responses, including dormancy release (Zhang et al. 2018). Therefore, integrating GA signaling into the bud dormancy model might be necessary to better account for the regulation of dormancy release. For example, it is possible that high GA levels around dormancy release play a role by overcoming the ABA-dependent growth inhibition. Interaction between GA and ABA pathways might also be critical in the response to environmental conditions during dormancy, including intertwined regulations of hormone biosynthesis (Shu et al. 2013, Yue et al. 2017). This was applied in a very innovative model for seed germination based on the endogenous hormone integration system (Topham et al. 2017). The hormonal balance between GA and ABA is regulated by endogenous and environmental signals towards the developmental switch that triggers termination of dormancy and germination. Accordingly, ON/OFF systems, like dormancy or flower initiation, can be modelled as developmental switches triggered in response to quantitative inputs after a threshold has been reached (Wilczek et al. 2009, Donohue et al. 2015, Bassel 2016). In our current model, we propose a first step for mechanistic modelling of dormancy onset and release based on expression data and ABA quantification. The next steps, in addition to the integration of GA signaling and its crosstalk with ABA, will be to provide information on temperature-mediated control of the regulatory cascades. Recent research led on Arabidopsis, allowed by high-throughput sequencing techniques, has hastened the pace for the incorporation of molecular data into phenology models (Satake et al. 2013, Kudoh 2016, Antoniou-Kourounioti et al. 2018, Nishio et al. 2020), thus opening new roads for perennial studies.

## DATA AVAILABILITY

RNA-seq data: Gene Expression Omnibus GSE130426

TPM data for the 81 genes analyzed in this work are available for all samples in the **supplementary datafile**

## SUPPLEMENTARY DATA

**Figure S1** Chilling and dormancy status during the treatments with exogenous hormones and antagonist on sweet cherry cultivar ‘Fertard‘

**Figure S2** Coverage of the mapped reads for each base of the sequences of ABA-related genes for the three sweet cherry cultivars

**Figure S3** Coverage of the mapped reads for each base of the sequences of GA-related genes for the three sweet cherry cultivars

**Figure S4** Effect of GAs, ABA and their inhibitor on the bud break percentage under forcing conditions

**Figure S5** Concentration ratio of ABA, ABA conjugates and GAs levels between ‘Regina’ and ‘Cristobalina‘

**Figure S6** ABA levels predictions for the best models using different sets of biosynthesis and catabolism genes

**Figure S7** Transcriptional dynamics of genes involved in ABA synthesis and degradation in the flower buds of the sweet cherry cultivar ‘Garnet‘

**Table S1** Description of the flower bud samples used for hormone quantification and RNA-seq and total number of mapped reads for RNA-seq data

**Table S2** Details of the genes related to ABA and GA pathways analyzed in the project

**Table S3** Differentially expressed genes

**Table S4** Parameters and residual sum of squares for the best models corresponding to the different gene sets tested to predict abscisic acid levels

**Supplementary Data File** Expression data in transcripts per million reads (TPM) for the genes analyzed in the project

## AKNOWLEDGEMENTS

The authors would like to thank INRAE and the Aquitaine Region for funding the doctoral fellowship to RB, and CMI-Groupe Roullier and ANRT for allowing the work conducted by NV. The authors warmly thank Teresa Barreneche, Hélène Christmann, Jacques Joly and Lydie Fouilhaux, for collecting the branches and collaborating on the phenotyping. The authors thank the INRAE‘s ‘*Prunus* Genetic Resources Center’ for preserving and managing the sweet cherry collections and the Fruit Experimental Unit of INRAE-Bordeaux (UEA) for growing the trees and managing the orchards. Figure 2 was modified from the original version published in Vimont et al. (2019), under copyright by BMC (Springer Nature Group).

## AUTHORS’ CONTRIBUTION

BW, SC, ED and PAW designed the original research. NV produced and analyzed the transcriptional data under the supervision of SC. RB conducted the analysis on exogenous application. LLD provided the sweet cherry genome on which ASLD performed the mapping and count analysis of RNA-seq data. AS and NV performed the phytohormones extraction and quantification. MA, FJ and JCY supervised the phytohormones analyses. NV and BW wrote the manuscript with the assistance of all the authors.

## REFERENCES

Ali N, Schwarzenberg A, Yvin JC, Hosseini SA (2018) Regulatory role of silicon in mediating differential stress tolerance responses in two contrasting tomato genotypes under osmotic stress. Front Plant Sci 9:1–16.

Anders S, Pyl PT, Huber W (2015) HTSeq-A Python framework to work with high-throughput sequencing data. Bioinformatics 31:166–169.

Antoniou-Kourounioti RL, Hepworth J, Heckmann A, Duncan S, Qüesta J, Rosa S, Säll T, Holm S, Dean C, Howard M (2018) Temperature Sensing Is Distributed throughout the Regulatory Network that Controls *FLC* Epigenetic Silencing in Vernalization. Cell Syst 7:643-655.e9.

Atkinson CJ, Brennan RM, Jones HG (2013) Declining chilling and its impact on temperate perennial crops. Environ Exp Bot 91:48–62.

Aubert D, Chevillard M, Dorne AM, Arlaud G, Herzog M (1998) Expression patterns of *GASA* genes in *Arabidopsis thaliana*: The *GASA4* gene is up-regulated by gibberellins in meristematic regions. Plant Mol Biol 36:871–883.

Bai S, Saito T, Sakamoto D, Ito A, Fujii H, Moriguchi T (2013) Transcriptome Analysis of Japanese Pear (*Pyrus pyrifolia* Nakai) Flower Buds Transitioning Through Endodormancy. Plant Cell Physiol 54:1132–1151.

Bassel GW (2016) To grow or not to grow. Trends Plant Sci 21:498–505.

Beauvieux R, Wenden B, Dirlewanger E (2018) Bud Dormancy in Perennial Fruit Tree Species : A Pivotal Role for Oxidative Cues. Front Plant Sci 9:1–13.

Bielenberg DG, Wang Y, Li Z, Zhebentyayeva T, Fan S, Reighard GL, Scorza R, Abbott AG (2008) Sequencing and annotation of the evergrowing locus in peach [*Prunus persica* (L.) Batsch] reveals a cluster of six MADS-box transcription factors as candidate genes for regulation of terminal bud formation. Tree Genet Genomes 4:495–507.

Bolger AM, Lohse M, Usadel B (2014) Trimmomatic: A flexible trimmer for Illumina sequence data. Bioinformatics 30:2114–2120.

Chmielewski F, Baldermann S, Götz K, Homann T, Gödeke K, Schumacher F, Huschek G, Rawel H (2018) Abscisic Acid Related Metabolites in Sweet Cherry Buds (*Prunus avium* L.). J Hortic 05.

Chmielewski F, Götz K, Homann T, Huschek G, Rawel H (2017) Identification of Endodormancy Release for Cherries (*Prunus Avium* L.) by Abscisic Acid and Sugars. J Hortic 04.

Chuine I, Bonhomme M, Legave J-M, García de Cortázar-Atauri I, Charrier G, Lacointe A, Améglio T (2016) Can phenological models predict tree phenology accurately in the future? The unrevealed hurdle of endodormancy break. Glob Chang Biol 22:3444–3460.

Chuine I, Régnière J (2017) Process-Based Models of Phenology for Plants and Animals. Annu Rev Ecol Evol Syst 48:159–82.

Cooke JEK, Eriksson ME, Junttila O (2012) The dynamic nature of bud dormancy in trees: environmental control and molecular mechanisms. Plant Cell Env 35:1707–1728.

Le Dantec L, Girollet N, Gouzy J, Sallet E, Carrère S, Fouché M, Quero-García J, Dirlewanger E (2020) Assembly and annotation of ‘Regina’ sweet cherry genome. V1. Portail Data INRAE. https://doi.org/10.15454/KEW474

Dobin A, Davis CA, Schlesinger F, Drenkow J, Zaleski C, Jha S, Batut P, Chaisson M, Gingeras TR (2013) STAR: Ultrafast universal RNA-seq aligner. Bioinformatics 29:15–21.

Donoho CWJ, Walker DR (1957) Effect of Gibberellic Acid on Breaking of Rest Period in Elberta Peach. Science (80-) 126:1178–1179.

Donohue K, Burghardt LT, Runcie D, Bradford KJ, Schmitt J (2015) Applying developmental threshold models to evolutionary ecology. Trends Ecol Evol 30:66–77.

Eriksson ME, Hoffman D, Kaduk M, Mauriat M, Moritz T (2015) Transgenic hybrid aspen trees with increased gibberellin (GA) concentrations suggest that GA acts in parallel with FLOWERING LOCUS T2 to control shoot elongation. New Phytol 205:1288–1295.

Fadón E, Herrero M, Rodrigo J (2015) Flower development in sweet cherry framed in the BBCH scale. Sci Hortic 192:141–147.

Fadón E, Rodrigo J, Herrero M (2018) Is there a specific stage to rest? Morphological changes in flower primordia in relation to endodormancy in sweet cherry (*Prunus avium* L.). Trees 32:1583–1594.

Finkelstein R (2013) Abscisic Acid Synthesis and Response. Arab B 11:e0166.

Fishman S, Erez A, Couvillon GA (1987a) The temperature dependence of dormancy breaking in plants: Mathematical analysis of a two-step model involving a cooperative transition. J Theor Biol 124:473–483.

Fishman S, Erez A, Couvillon GA (1987b) The temperature dependence of dormancy breaking in plants: Computer simulation of processes studied under controlled temperatures. J Theor Biol 126:309–321.

Gagne JM, Downes BP, Shiu S-H, Durski AM, Vierstra RD (2002) The F-box subunit of the SCF E3 complex is encoded by a diverse superfamily of genes in Arabidopsis. Proc Natl Acad Sci 99:11519–11524.

Haddad C, Arkoun M, Jamois F, Schwarzenberg A, Yvin JC, Etienne P, Laîné P (2018) Silicon promotes growth of Brassica napus L. And delays leaf senescence induced by nitrogen starvation. Front Plant Sci 9:1–13.

Hänninen H (1990) Modelling bud dormancy release in trees from cool and temperate regions. Acta For Fenn 213:1–47.

Hao X, Chao WS, Yang Y, Horvath DP (2015) Coordinated expression of *FLOWERING LOCUS T* and *DORMANCY ASSOCIATED MADS-BOX-like* genes in leafy spurge. PLoS One 10:1–18.

Heide OM, Prestrud AK (2005) Low temperature, but not photoperiod, controls growth cessation and dormancy induction and release in apple and pear. Tree Physiol 25:109–114.

Hoad G V (1983) Hormonal regulation of fruit-bud formation in fruit trees. Acta Hortic 149:13–24.

Horvath DP (2009) Common mechanisms regulate flowering and dormancy. Plant Sci 177:523–531.

Howe GT, Horvath DP, Dharmawardhana P, Priest HD, Mockler TC, Strauss SH (2015) Extensive Transcriptome Changes During Natural Onset and Release of Vegetative Bud Dormancy in Populus. Front Plant Sci 6

Huerta-cepas J, Forslund K, Coelho LP, Szklarczyk D, Jensen LJ, von Mering C, Bork P (2017) Fast Genome-Wide Functional Annotation through Orthology Assignment by eggNOG-Mapper. Mol Biol Evol 34:2115–2122.

Huerta-cepas J, Szklarczyk D, Heller D, Hernandez-Plaza A, Forslund SK, Cook H, Mende DR, Letunic I, Rattei T, Jensen LJ, von Mering C, Bork P (2019) eggNOG 5.0: a hierarchical, functionally and phylogenetically annotated orthology resource based on 5090 organisms and 2502 viruses. Nucleic Acids Res 47:309–314.

Ito A, Tuan PA, Saito T, Bai S, Kita M, Moriguchi T (2019) Changes in Phytohormone Content and Associated Gene Expression Throughout the Stages of Pear (*Pyrus pyrifolia* Nakai) Dormancy. Tree Physiol tpz101

Jiménez S, Reighard GL, Bielenberg DG (2010) Gene expression of DAM5 and DAM6 is suppressed by chilling temperatures and inversely correlated with bud break rate. Plant Mol Biol 73:157–67. http://www.ncbi.nlm.nih.gov/pubmed/20143130 (30 September 2013, date last accessed).

Jochner S, Caffarra A, Menzel A (2013) Can spatial data substitute temporal data in phenological modelling? A survey using birch flowering. Tree Physiol 33:1256–1268.

Junttila O, Jensen E (1988) Gibberellins and photoperiodic control of shoot elongation in Salix. Physiol Plant 74:371–376.

El Kayal W, Allen CCG, Ju CJT, Adams E, King-Jones S, Zaharia LI, Abrams SR, Cooke JEK (2011) Molecular events of apical bud formation in white spruce, Picea glauca. Plant Cell Environ 34:480–500.

Khalil-Ur-Rehman M, Sun L, Li CX, Faheem M, Wang W, Tao JM (2017) Comparative RNA-seq based transcriptomic analysis of bud dormancy in grape. BMC Plant Biol 17:1–11.

Kudoh H (2016) Molecular phenology in plants: In natura systems biology for the comprehensive understanding of seasonal responses under natural environments. New Phytol 210:399–412.

de la Fuente L, Conesa A, Lloret A, Badenes ML, Ríos G (2015) Genome-wide changes in histone H3 lysine 27 trimethylation associated with bud dormancy release in peach. Tree Genet Genomes 11:45.

Lakkis S, Trotel-Aziz P, Rabenoelina F, Schwarzenberg A, Nguema-Ona E, Clément C, Aziz A (2019) Strengthening Grapevine Resistance by Pseudomonas fluorescens PTA-CT2 Relies on Distinct Defense Pathways in Susceptible and Partially Resistant Genotypes to Downy Mildew and Gray Mold Diseases. Front Plant Sci 10:1–18.

Lang G, Early J, Martin G, Darnell R (1987) Endo-, para-, and ecodormancy: physiological terminology and classification for dormancy research. Hort Sci 22:371–377.

Leida C, Conejero A, Arbona V, Gómez-Cadenas A, Llácer G, Badenes ML, Ríos G (2012) Chilling-dependent release of seed and bud dormancy in peach associates to common changes in gene expression. PLoS One 7:e35777.

Leida C, Conesa A, Llácer G, Badenes ML, Ríos G (2012) Histone modifications and expression of *DAM6* gene in peach are modulated during bud dormancy release in a cultivar-dependent manner. New Phytol 193:67–80.

Li J, Xu Y, Niu Q, He L, Teng Y, Bai S (2018) Abscisic acid (ABA) promotes the induction and maintenance of pear (*Pyrus pyrifolia* white pear group) flower bud endodormancy. Int J Mol Sci 19

Liu J, Sherif SM (2019) Hormonal Orchestration of Bud Dormancy Cycle in Deciduous Woody Perennials. Front Plant Sci 10:1–21.

Love MI, Huber W, Anders S (2014) Moderated estimation of fold change and dispersion for RNA-seq data with DESeq2. Genome Biol 15:1–21.

Nambara E, Marion-Poll A (2005) Abscisic Acid Biosynthesis and Catabolism. Annu Rev Plant Biol 56:165–185.

Nishio H, Buzas DM, Nagano AJ, Iwayama K, Ushio M, Kudoh H (2020) Repressive chromatin modification underpins the long-term expression trend of a perennial flowering gene in nature. Nat Commun 11.

Niu Q, Li J, Cai D, Qian M, Jia H, Bai S, Hussain S, Liu G, Teng Y, Zheng X (2015) Dormancy-associated MADS-box genes and microRNAs jointly control dormancy transition in pear (*Pyrus pyrifolia* white pear group) flower bud. J Exp Bot 67:erv454.

Olsen JE (2010) Light and temperature sensing and signaling in induction of bud dormancy in woody plants. Plant Mol Biol 73:37–47.

Ophir R, Pang X, Halaly T, Venkateswari J, Lavee S, Galbraith D, Or E (2009) Gene-expression profiling of grape bud response to two alternative dormancy-release stimuli expose possible links between impaired mitochondrial activity, hypoxia, ethylene-ABA interplay and cell enlargement. Plant Mol Biol 71:403–423.

Or E, Belausov E, Popilevsky I, Ben Tal Y (2000) Changes in endogenous ABA level in relation to the dormancy cycle in grapevines grown in a hot climate. J Hortic Sci Biotechnol 75:190–194.

Petterle A, Karlberg A, Bhalerao RP (2013) Daylength mediated control of seasonal growth patterns in perennial trees. Curr Opin Plant Biol 16:301–306.

Powell LE (1987) The hormonal control of bud and seed dormancy in woody plants. In: Davies P (ed) Plant Hormones and Their Role in Plant Growth and Development. Martinus Nijhoff Publishers, Dordrecht, pp 539–552.

Priest DM, Ambrose SJ, Vaistij FE, Elias L, Higgins GS, Ross ARS, Abrams SR, Bowles DJ (2006) Use of the glucosyltransferase UGT71B6 to disturb abscisic acid homeostasis in Arabidopsis thaliana. Plant J 46:492–502.

Rinne PLH, Welling A, Vahala J, Ripel L, Ruonala R, Kangasjarvi J, van der Schoot C (2011) Chilling of dormant buds hyperinduces FLOWERING LOCUS T and recruits GA-inducible 1,3-beta-glucanases to reopen signal conduits and release dormancy in Populus. Plant Cell 23:130–146.

Ríos G, Leida C, Conejero A, Badenes ML (2014) Epigenetic regulation of bud dormancy events in perennial plants. Front Plant Sci 5:247.

Rodríguez-Gacio MDC, Matilla-Vázquez M a, Matilla AJ (2009) Seed dormancy and ABA signaling: the breakthrough goes on. Plant Signal Behav 4:1035–1049.

Rodriguez A, Sherman W, Scorza R, Wisniewski M, Okie WR (1994) ‘Evergreen’ peach, its inheritance and dormant behavior. J Am Soc hort Sci 119:789–792.

Rohde A, Bhalerao RP (2007) Plant dormancy in the perennial context. Trends Plant Sci 12:217–223.

Rohde A, Prinsen E, Rycke R De, Engler G, Montagu M Van, Boerjan W (2002) PtABI3 Impinges on the Growth and Differentiation of Embryonic Leaves during Bud Set in Poplar. Plant Cell 14:1885–1901.

Rohde A, Storme V, Jorge V, Gaudet M, Vitacolonna N, Fabbrini F, Ruttink T, Zaina G, Marron N, Dillen S, Steenackers M, Sabatti M, Morgante M, Boerjan W, Bastien C (2011) Bud set in poplar--genetic dissection of a complex trait in natural and hybrid populations. New Phytol 189:106–121.

Roxrud I, Lid SE, Fletcher JC, Schmidt EDL, Opsahl-Sorteberg HG (2007) GASA4, one of the 14-member Arabidopsis GASA family of small polypeptides, regulates flowering and seed development. Plant Cell Physiol 48:471–483.

Ruttink T, Arend M, Morreel K, Storme V, Rombauts S, Fromm J, Bhalerao RP, Boerjan W, Rohde A (2007) A molecular timetable for apical bud formation and dormancy induction in poplar. Plant Cell 19:2370–2390.

Satake A, Kawagoe T, Saburi Y, Chiba Y, Sakurai G, Kudoh H (2013) Forecasting flowering phenology under climate warming by modelling the regulatory dynamics of flowering-time genes. Nat Commun 4:2303.

Schwacke R, Ponce-soto GY, Krause K, Bolger AM, Arsova B, Hallab A, Gruden K, Stitt M, Bolger ME, Usadel B (2019) MapMan4: A Refined Protein Classification and Annotation Framework Applicable to Multi-Omics Data Analysis. Mol Plant 12:879–892.

Shafer N, Monson WG (1958) The Role og Gibberellic Acid in Overcoming Bud Dormancy in Perennial Weeds. I. Leafy Spurge (Euphorbia esulta L.) and Ironweed (Vernonia Baldwini Torr.). Weeds 6:172–178.

Shu K, Zhang H, Wang S, Chen M, Wu Y, Tang S, Liu C, Feng Y, Cao X, Xie Q (2013) ABI4 Regulates Primary Seed Dormancy by Regulating the Biogenesis of Abscisic Acid and Gibberellins in Arabidopsis. PLoS Genet 9

Shu K, Zhou W, Yang W (2018) APETALA 2-domain-containing transcription factors: focusing on abscisic acid and gibberellins antagonism. New Phytol 217:977–983.

Singh RK, Maurya JP, Azeez A, Miskolczi P, Tylewicz S, Stojkovic K, Delhomme N, Busov V, Bhalerao RP (2018) A genetic network mediating the control of bud break in hybrid aspen. Nat Commun 9

Singh RK, Miskolczi P, Maurya JP, Bhalerao RP (2019) A Tree Ortholog of SHORT VEGETATIVE PHASE Floral Repressor Mediates Photoperiodic Control of Bud Dormancy. Curr Biol 29:128-133.e2.

Singh RK, Svystun T, AlDahmash B, Jönsson AM, Bhalerao RP (2016) Photoperiod- and temperature-mediated control of growth cessation and dormancy in trees: A molecular perspective. New Phytol 213:511–524.

Topham AT, Taylor RE, Yan D, Nambara E, Johnston IG, Bassel GW (2017) Temperature variability is integrated by a spatially embedded decision-making center to break dormancy in Arabidopsis seeds. Proc Natl Acad Sci 114:6629–6634.

Tuan PA, Bai S, Saito T, Ito A, Moriguchi T (2017) Dormancy-Associated MADS-Box (DAM) and the Abscisic Acid Pathway Regulate Pear Endodormancy Through a Feedback Mechanism. Plant Cell Physiol 58:1378–1390.

Tylewicz S, Petterle A, Marttila S, Miskolczi P, Azeez A, Singh RK, Immanen J, Mähler N, Hvidsten TR, Eklund DM, Bowman JL, Helariutta Y, Bhalerao RP (2018) Photoperiodic control of seasonal growth is mediated by ABA acting on cell-cell communication. Science (80-) 360:212–215.

Ueguchi-Tanaka M, Nakajima M, Katoh E, Ohmiya H, Asano K, Saji S, Hongyu X, Ashikari M, Kitano H, Yamaguchi I, Matsuoka M (2007) Molecular Interactions of a Soluble Gibberellin Receptor, GID1, with a Rice DELLA Protein, SLR1, and Gibberellin. Plant Cell Online 19:2140–2155.

Ueno S, Klopp C, Leplé JC, Derory J, Noirot C, Léger V, Prince E, Kremer A, Plomion C, Le Provost G (2013) Transcriptional profiling of bud dormancy induction and release in oak by next-generation sequencing. BMC Genomics 14:236.

Vimont N, Fouché M, Campoy JA, Tong M, Arkoun M, Yvin J-C, Wigge PA, Dirlewanger E, Cortijo S, Wenden B (2019) From bud formation to flowering: transcriptomic state defines the cherry developmental phases of sweet cherry bud dormancy. BMC Genomics 20:974. doi:

Vitasse Y, François C, Delpierre N, Dufrêne E, Kremer A, Chuine I, Delzon S (2011) Assessing the effects of climate change on the phenology of European temperate trees. Agric For Meteorol 151:969–980.

Wagner GP, Kin K, Lynch VJ (2012) Measurement of mRNA abundance using RNA-seq data: RPKM measure is inconsistent among samples. Theory Biosci 131:281–285.

Wang D, Gao Z, Du P, Xiao W, Tan Q, Chen X, Li L, Gao D (2016) Expression of ABA Metabolism-Related Genes Suggests Similarities and Differences Between Seed Dormancy and Bud Dormancy of Peach (Prunus persica). Front Plant Sci 6:1–17.

Weinberger J (1950) Chilling requirements of peach varieties. Proc Am Soc Hortic Sci 56:122–128.

Wen LH, Zhong WJ, Huo XM, Zhuang WB, Ni ZJ, Gao ZH (2016) Expression analysis of ABA- and GA-related genes during four stages of bud dormancy in Japanese apricot (*Prunus mume* Sieb. et Zucc). J Hortic Sci Biotechnol 91:362–369.

Weng JK, Ye M, Li B, Noel JP (2016) Co-evolution of Hormone Metabolism and Signaling Networks Expands Plant Adaptive Plasticity. Cell 166:881–893.

Wilczek AM, Roe JL, Knapp MC, Cooper MD, Lopez-Gallego C, Martin LJ, Muir CD, Sim S, Walker A, Anderson J, Egan JF, Moyers BT, Petipas R, Giakountis A, Charbit E, Coupland G, Welch SM, Schmitt J, Franklin Egan J, Moyers BT, Petipas R, Giakountis A, Charbit E, Coupland G, Welch SM, Schmitt J (2009) Effects of genetic perturbation on seasonal life history plasticity. Science (80) 323:930–935.

Xu ZJ, Nakajima M, Suzuki Y, Yamaguchi I (2002) Cloning and Characterization of the Abscisic Acid-Specific Glucosyltransferase Gene from Adzuki Bean Seedlings. Plant Physiol 129:1285–1295.

Yamaguchi S (2008) Gibberellin Metabolism and its Regulation. Annu Rev Plant Biol 59:225–251.

Yamane H, Wada M, Honda C, Matsuura T, Ikeda Y, Hirayama T, Osako Y, Gao-Takai M, Kojima M, Sakakibara H, Tao R (2019) Overexpression of Prunus DAM6 inhibits growth, represses bud break competency of dormant buds and delays bud outgrowth in apple plants. PLoS One 14:1–24.

Yang Q, Niu Q, Tang Y, Ma Y, Yan X, Li J, Tian J, Bai S, Teng Y (2019) PpyGAST1 is potentially involved in bud dormancy release by integrating the GA biosynthesis and ABA signaling in ‘Suli’ pear (Pyrus pyrifolia White Pear Group). Environ Exp Bot 162:302–312.

Yang Q, Yang B, Li J, Wang Y, Tao R, Yang F, Wu X, Yan X, Ahmad M, Shen J, Bai S, Teng Y (2020) ABA-responsive ABRE-BINDING FACTOR3 activates DAM3 expression to promote bud dormancy in Asian pear. Plant Cell Environ 43:1360–1375.

Yu J, Conrad AO, Decroocq V, Zhebentyayeva T, Williams DE, Bennett D, Roch G, Audergon JM, Dardick C, Liu Z, Abbott AG, Staton ME (2020) Distinctive Gene Expression Patterns Define Endodormancy to Ecodormancy Transition in Apricot and Peach. Front Plant Sci 11:1–24.

Yue C, Cao H, Hao X, Zeng J, Qian W, Guo Y, Ye N, Yang Y, Wang X (2017) Differential expression of gibberellin- and abscisic acid-related genes implies their roles in the bud activity-dormancy transition of tea plants. Plant Cell Rep 0:0.

Zentella R, Zhang Z-L, Park M, Thomas SG, Endo A, Murase K, Fleet CM, Jikumaru Y, Nambara E, Kamiya Y, Sun T (2007) Global Analysis of DELLA Direct Targets in Early Gibberellin Signaling in *Arabidopsis*. Plant Cell 19:3037–3057.

Zhang X, An L, Nguyen TH, Liang H, Wang R, Liu X, Li T, Qi Y, Yu F (2015) The cloning and functional characterization of peach *CONSTANS* and *FLOWERING LOCUS T* homologous genes PpCO and PpFT. PLoS One 10:1–16.

Zhang Z, Zhuo X, Zhao K, Zheng T, Han Y, Yuan C, Zhang Q (2018) Transcriptome Profiles Reveal the Crucial Roles of Hormone and Sugar in the Bud Dormancy of *Prunus mume*. Sci Rep 8:1–15.

Zhao K, Zhou Y, Ahmad S, Yong X, Xie X, Han Y, Li Y, Sun L, Zhang Q (2018) PmCBFs synthetically affect PmDAM6 by alternative promoter binding and protein complexes towards the dormancy of bud for *Prunus mume*. Sci Rep 8:4527.

Zheng C, Acheampong AK, Shi Z, Halaly T, Kamiya Y, Ophir R, Galbraith DW, Or E (2018) Distinct gibberellin functions during and after grapevine bud dormancy release. J Exp Bot 69:1635–1648.

Zheng C, Acheampong AK, Shi Z, Mugzech A, Halaly-Basha T, Sun Y, Colova V, Mosquna A, Ophir R, Galbraith DW, Or E (2018) Abscisic Acid Catabolism Enhances Dormancy Release of Grapevine Buds. Plant Cell Environ 41:2490–2503.

Zheng C, Halaly T, Acheampong AK, Takebayashi Y, Jikumaru Y, Kamiya Y, Or E (2015) Abscisic acid (ABA) regulates grape bud dormancy, and dormancy release stimuli may act through modification of ABA metabolism. J Exp Bot 66:1527–1542.

Zhong W, Gao Z, Zhuang W, Shi T, Zhang Z, Ni Z (2013) Genome-wide expression profiles of seasonal bud dormancy at four critical stages in Japanese apricot. Plant Mol Biol 83:247–64.

Zhong C, Xu H, Ye S, Wang S, Li L, Zhang S, Wang X (2015) AtGASA6 Serves as an Integrator of Gibberellin-, Abscisic Acid- and Glucose-Signaling during Seed Germination in Arabidopsis. Plant Physiol 169:pp.00858.2015.

Zhu Y, Li Y, Xin D, Chen W, Shao X, Wang Y, Guo W (2015) RNA-Seq-based transcriptome analysis of dormant flower buds of Chinese cherry (*Prunus pseudocerasus*). Gene 555:362–376.

Zhuang W, Gao Z, Wang L, Zhong W, Ni Z, Zhang Z (2013) Comparative proteomic and transcriptomic approaches to address the active role of GA4 in Japanese apricot flower bud dormancy release. J Exp Bot 64:4953–4966.

