## Supplementary Figures for "Fine tuning of hormonal signaling is linked to dormancy status in sweet cherry flower buds"

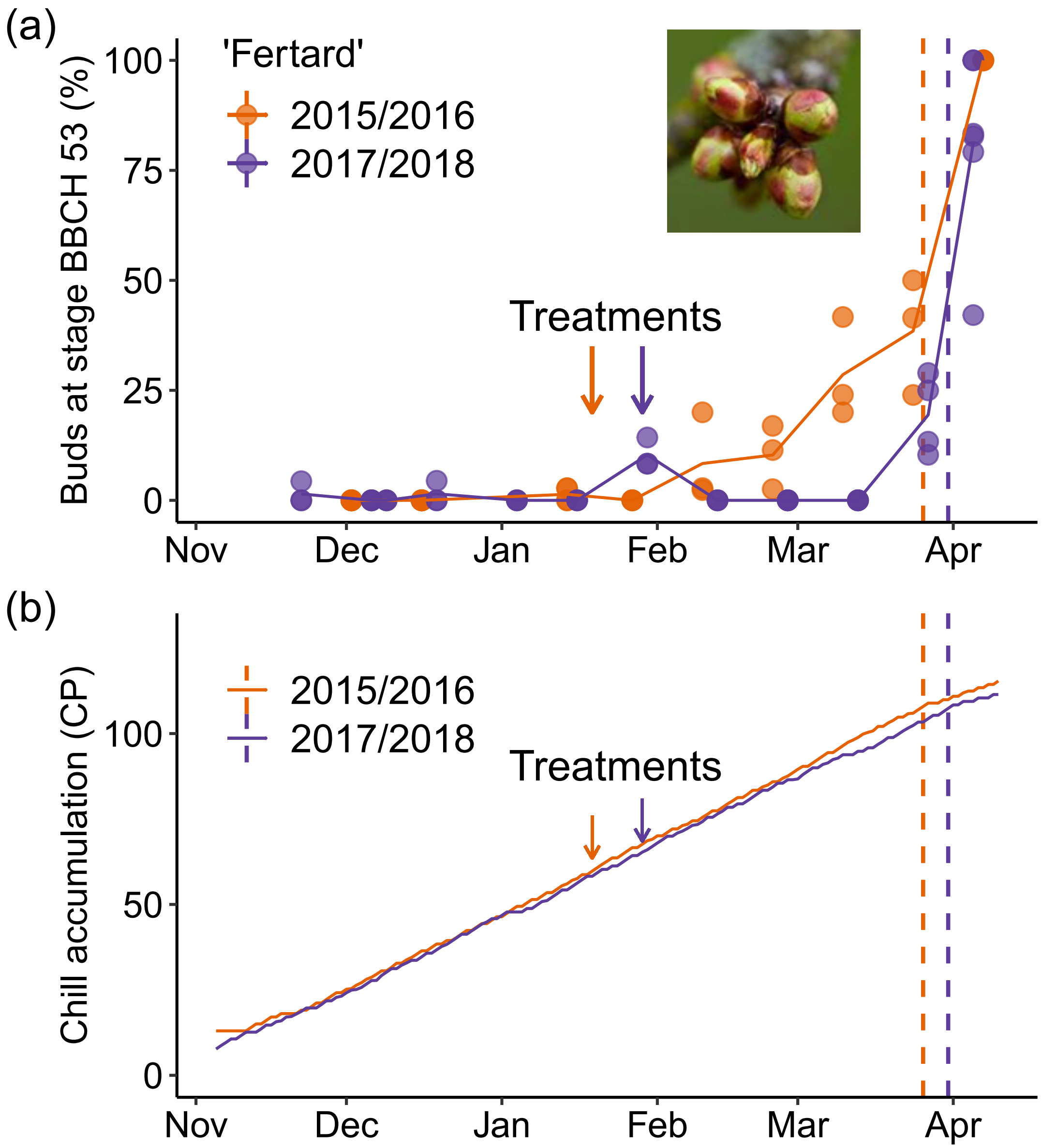


**Figure S1. Chilling and dormancy status during the treatments with exogenous hormones and antagonist on sweet cherry cultivar 'Fertard'**

(a) Evaluation of bud break percentage under forcing conditions was conducted on branches carrying flower buds. Branches were randomly sampled from three trees and placed under forcing conditions (25°C, 16h light, 8h dark). After ten days, the percentage of bud break, i.e. flower buds at BBCH 53 stage (see picture) was recorded. The date of dormancy release (dashed lines) was estimated when at least 50% of the flower buds were at the BBCH 53 stage or higher. (b) Chill accumulation calculated in chill portions (CP, as estimated by the Dynamic model) was calculated for the two treatment seasons. Arrows indicate the date at which branches were sampled, treated with exogenous hormones and antagonists and then placed under forcing conditions.

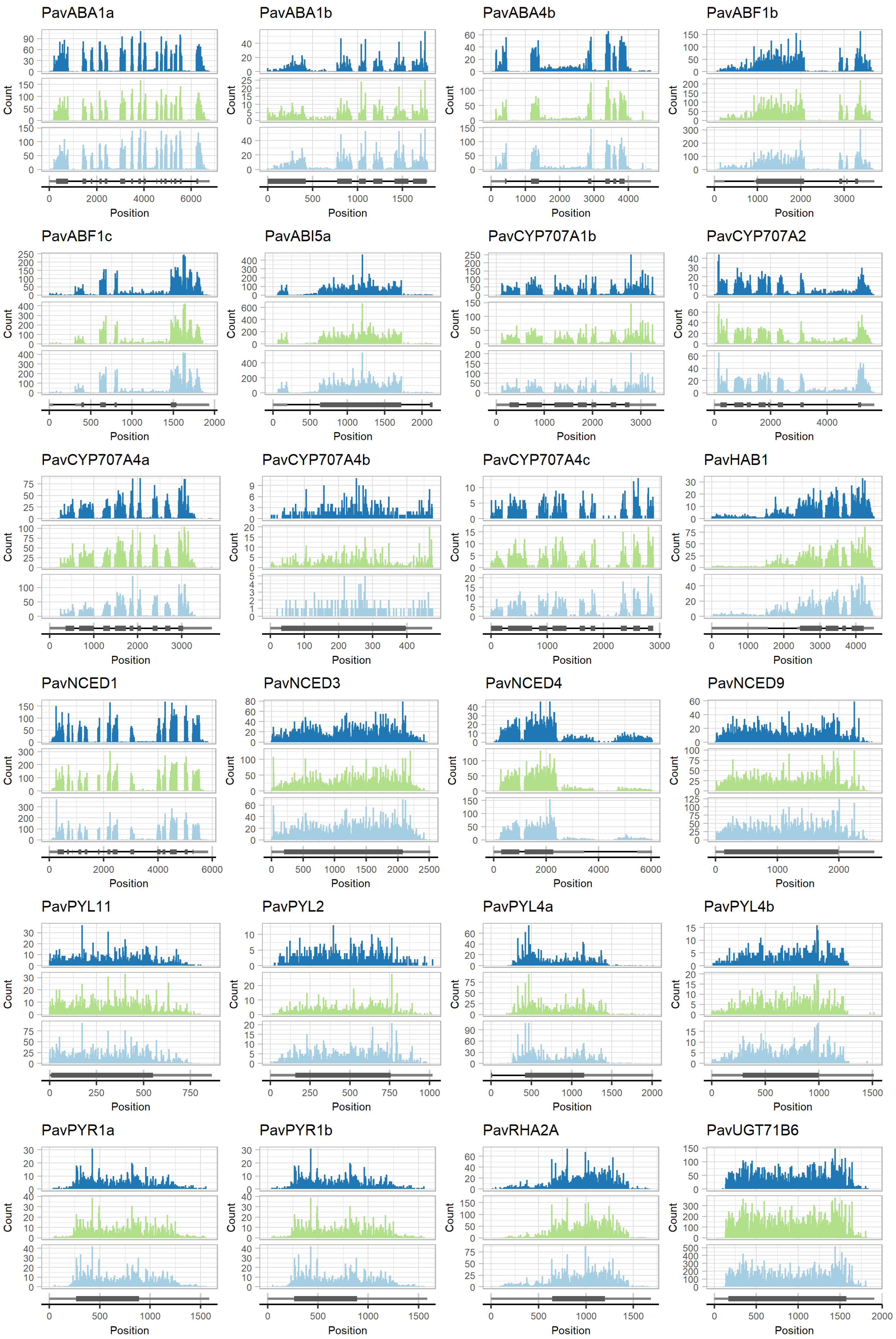


**Figure S2. Coverage of the mapped reads for each base of the sequences of ABA-related genes for the three sweet cherry cultivars** Mapped reads from all samples were used for the count. Upper panel (dark blue): 'Regina', middle panel (green): 'Garnet', lower panel (light blue): 'Cristobalina'. Dark rectangles correspond to the predicted exons of the peach gene and light grey rectangles represent the 3' and 5' UTR sequences. ABA: Abscisic acid; ABA1/4: ABA DEFICIENT 1/4; ABF1: ABSCISIC ACID RESPONSIVE ELEMENTS-BINDING PROTEIN; ABI5: ABA INSENSITIVE 5; HAB: homology to ABI2; NCED: 9-cis epoxycarotenoid dioxygenase; PYR: PYRABACTIN RESISTANCE; PYL: PYR-like; RHA2A: RING;-H2 A; UGT: UDP-GLYCOSYLTRANSFERASE.

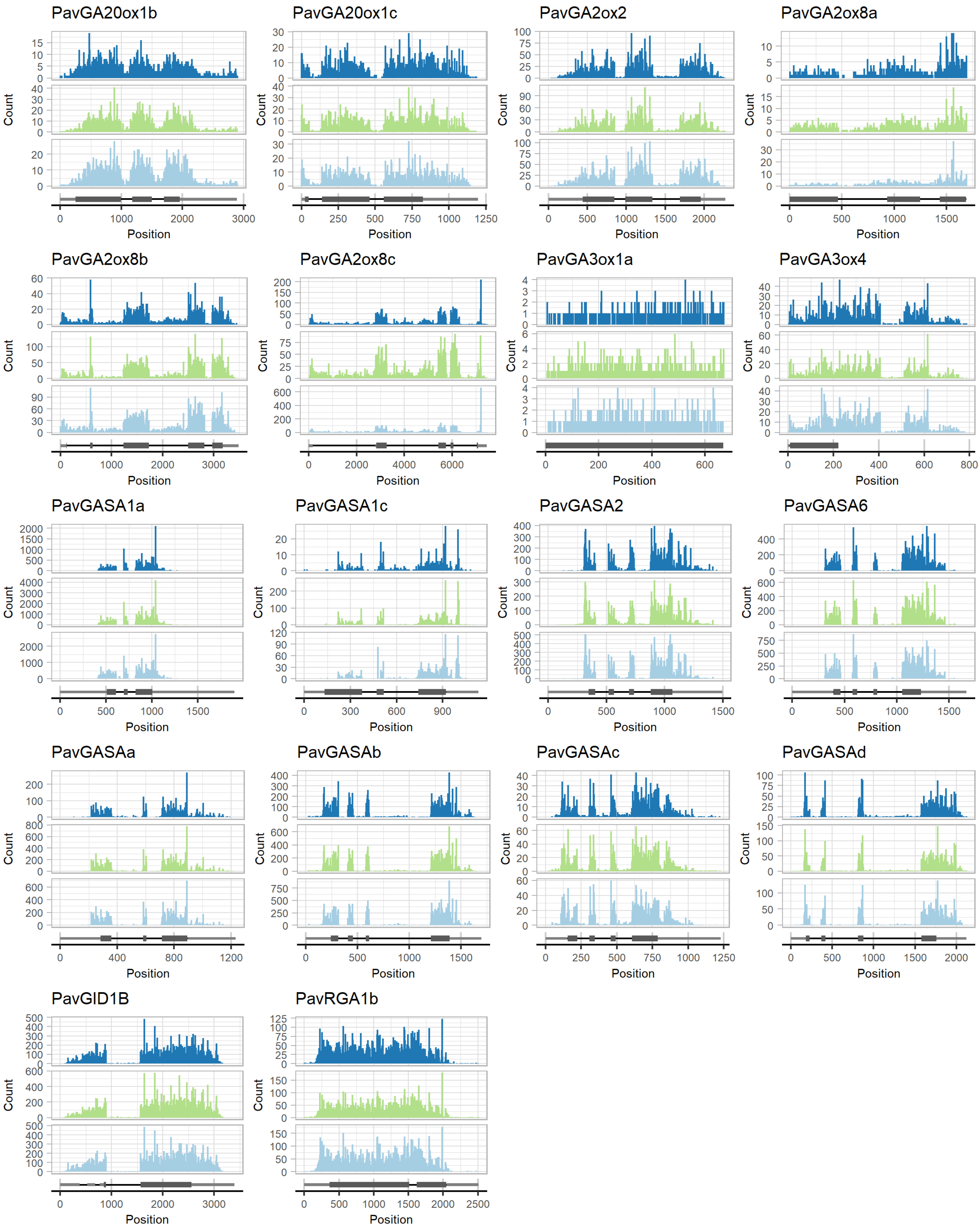


**Figure S3. Coverage of the mapped reads for each base of the sequences of GA-related genes for the three sweet cherry cultivars** Mapped reads from all samples were used for the count. Upper panel (dark blue): 'Regina', middle panel (green): 'Garnet', lower panel (light blue): 'Cristobalina'. Dark rectangles correspond to the predicted exons of the peach gene and light grey rectangles represent the 3' and 5' UTR sequences. GA: Gibberellic acid; GA20ox: GA 20-oxidases, GA3ox: GA 3-oxidases; GA2ox: GA 2-oxidases; GID: GA INSENSITIVE DWARF; GASA: GA Stimulated Arabidopsis; RGA: REPRESSOR OF GA.

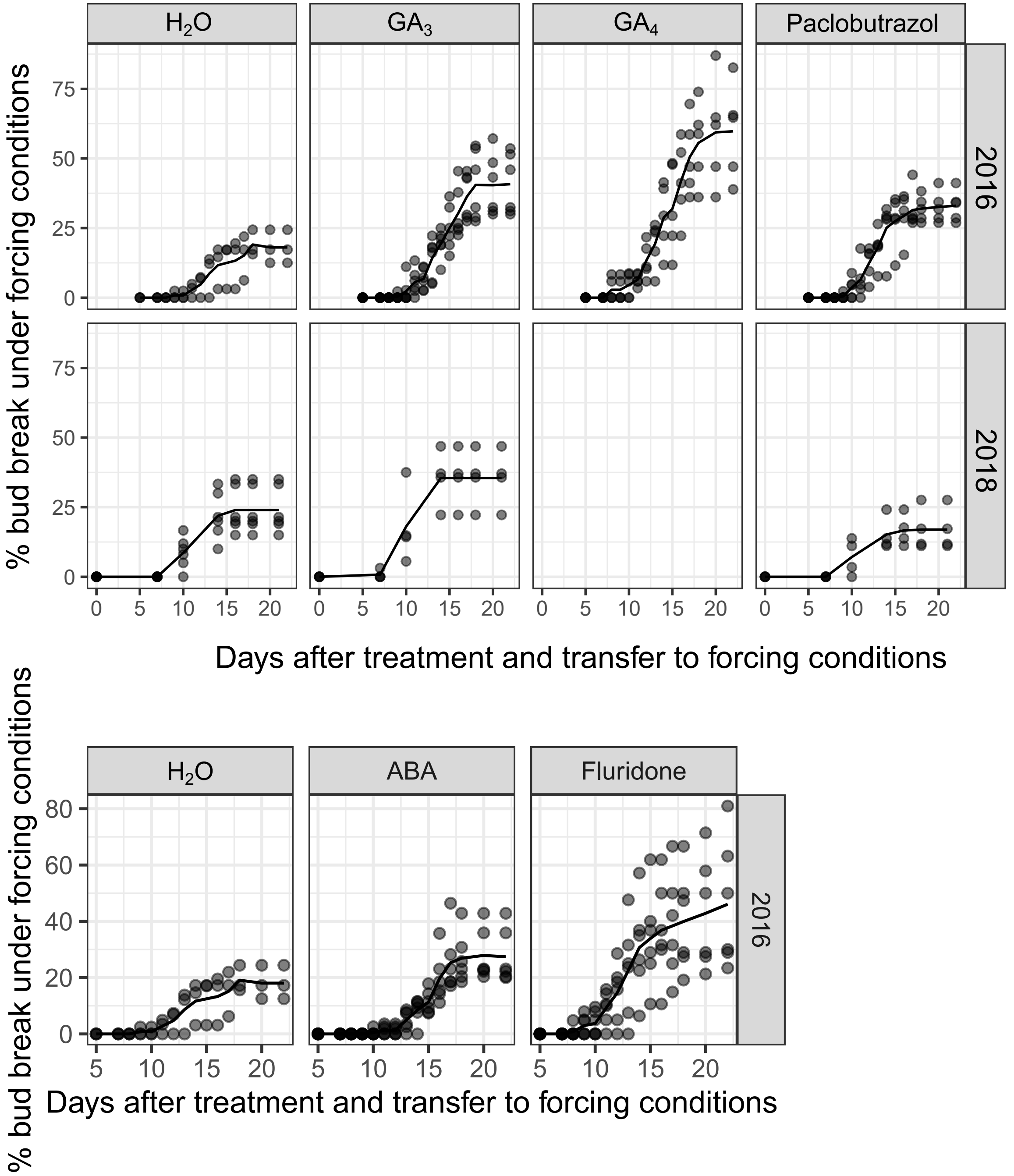


**Figure S4. Effect of different GAs, ABA and their inhibitor on the bud break percentage under forcing conditions**

Sweet cherry branches carrying dormant flower buds were treated with 5µM GA3, 5 µM GA4, 300 µM paclobutrazol (GA pathway inhibitor, 400 µM ABA and 5 µM fluridone (ABA pathway inhibitor) and transferred under forcing conditions (25°C, 60-70% humidity, 16 hours light). The percentage of flower bud break was recorded three times a week. The line corresponds to the average of biological replicates (3 to 5 depending on the treatment). ABA: Abscisic acid; GA: Gibberellic acid.

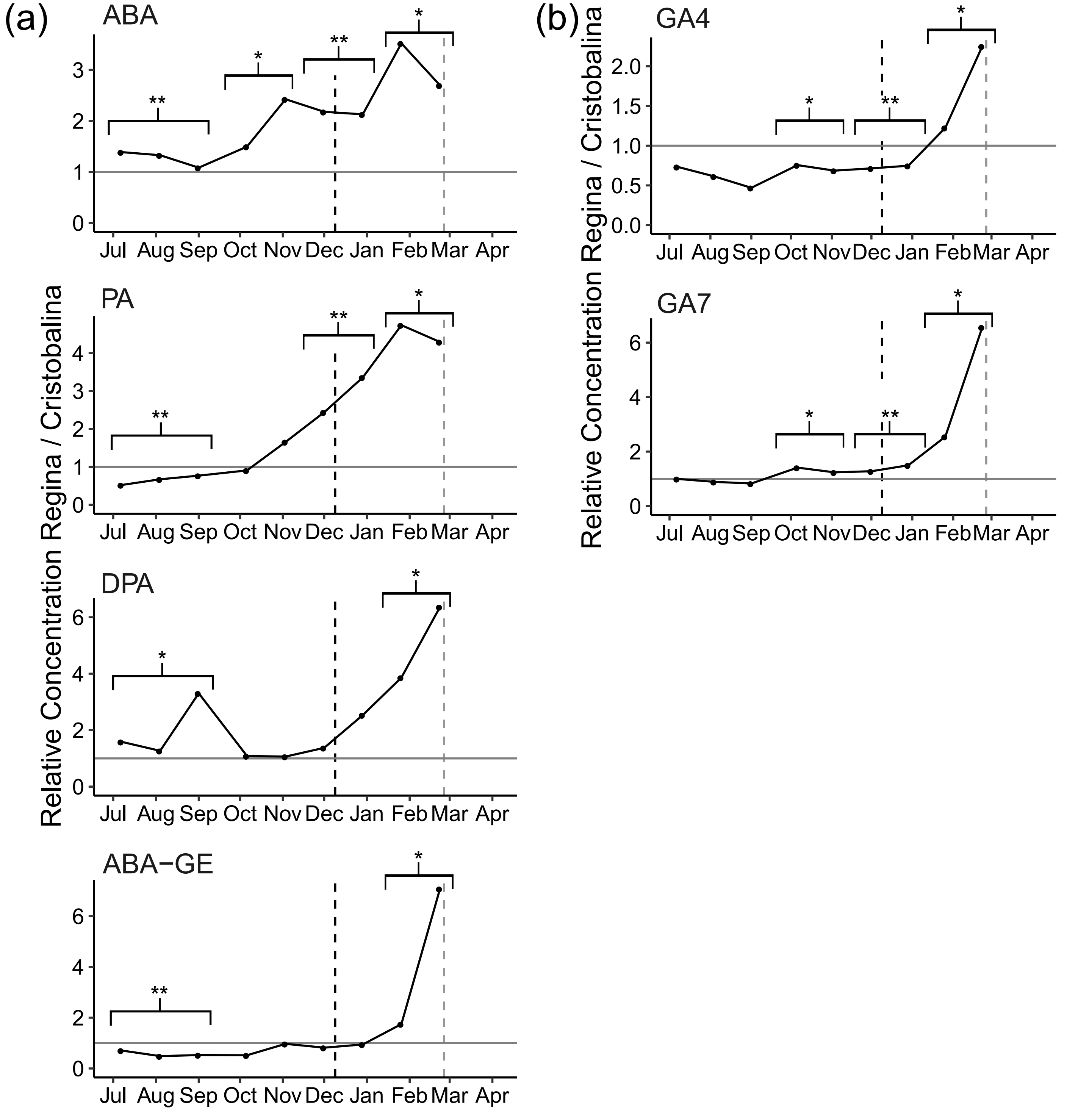


**Figure S5. Concentration ratio of ABA, ABA conjugates and GAs levels between 'Regina' and 'Cristobalina'**

The relative concentrations for (a) ABA and ABA conjugates and (b) GAs were calculated as the concentration ratio between 'Regina' and 'Cristobalina' for each common sampling date (see Figures 4 and 6 for the absolute values). Dotted lines represent the dormancy release date estimated for the early flowering cultivar 'Cristobalina' (black) and the late flowering cultivar 'Regina' (grey). Asterisks indicate significantly different levels between the two cultivars for grouped dates (1: July-September; 2: October-November; 3: December; 4: January-February - Kruskal-Wallis test, (*): *p-value* < 0.05; (**): *p-value* < 0.01; (***): *p-value* < 0.005). ABA: Abscisic acid; PA: phaseic acid; DPA: dihydrophaseic acid; ABA-GE: ABA-glucose ester; GA: gibberellic acid.

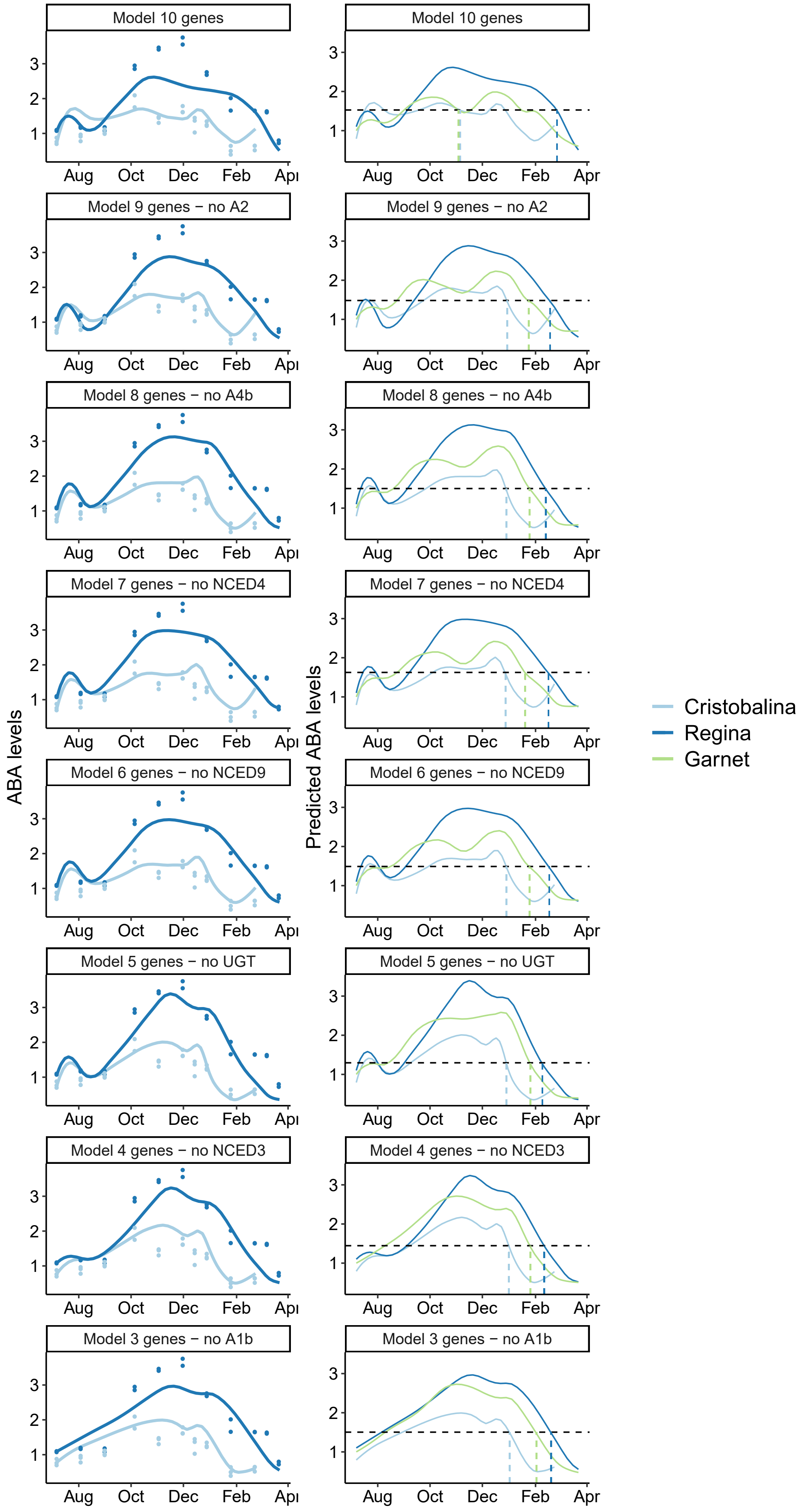


**Figure S6. ABA levels predictions for the best models using different sets of biosynthesis and catabolism genes**
(Left panel) Simulated content of ABA in 'Regina' and 'Cristobalina' using the best models for 3, 4, 5, 6, 7, 8, 9 and 10 genes. Lines represent the simulated ABA levels and circles represent the experimental data. (Right panel) Simulated levels of ABA for cultivars 'Cristobalina', 'Regina' and 'Garnet'. An ABA threshold (black dash line) was estimated for dormancy breaking (colored dash lines). Details on the models can be found in Table S4.

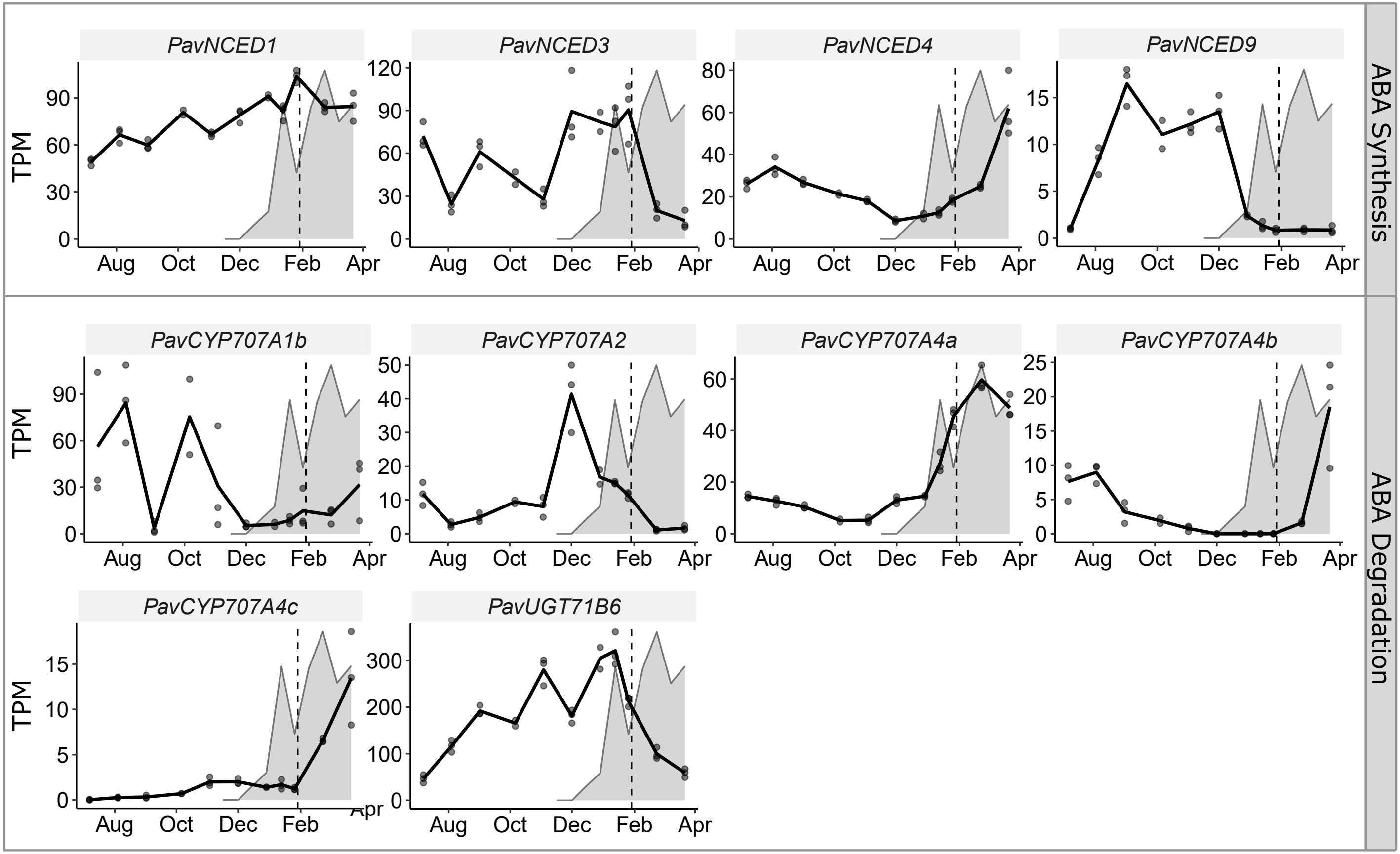


**Figure S7. Transcriptional dynamics of genes associated with ABA synthesis and degradation in the flower buds of the sweet cherry cultivar 'Garnet'**

Expression of specific genes involved in ABA biosynthesis and degradation are represented in TPM (transcripts per million reads). Dots indicate the data for the biological replicates (see Table S1 for details). Background areas correspond to the dormancy depth evaluated as the percentage of bud break under forcing conditions (see Fig. 2). Dotted lines represent dormancy release. ABA: Abscisic acid; NCED: 9-cis epoxycarotenoid dioxygenase; UGT: UDP-GLYCOSYLTRANSFERASE.
